# A novel fracture lattice in spiny mouse skin facilitates tissue autotomy and regeneration

**DOI:** 10.64898/2026.03.23.713756

**Authors:** Daeryeok Ko, Yeong Chan Ryu, Jae-Hoon Choi, Eunu Kim, Hyunji Cha, Soyun Joo, Seunghwan Ryu, Hyemin Ryu, Sungwook Shim, Jiyeon Lee, Seulki You, Jiwon Lim, Jie Tong, Catherine P. Lu, Sooil Chang, Ji Ae Kim, Ji Won Oh, Ann M. Clemens, Ashley W. Seifert, Seungbum Hong, Haeshin Lee, Gi-Dong Sim, Hanseul Yang

## Abstract

Autotomy is a unique phenotype whereby an animal sheds a body part to escape predation^1–3^. The timing and location of autotomy are tightly regulated by preformed planes of weakness (aka fracture planes) which facilitate tissue loss. While autotomy is often followed by regeneration, these phenotypes are rarely reported in mammals^4–9^. A notable exception are spiny mice (*Acomys*) which exhibit skin autotomy and more remarkably, complete tissue regeneration^10–14^. Presently, mechanisms underlying autotomy and complete regeneration in *Acomys* skin remain elusive. Here, we report the discovery of a honeycomb-like fracture lattice in *Acomys* skin whose design directs tissue destruction but also facilitates regenerative healing. Unlike the single continuous surface of a fracture plane, this fracture lattice consists of a three-dimensional array of hexagonal units whose boundaries guide tissue breakage. Moreover, we identify collagen VI as the main constituent of the fracture lattice and find that it is distinctly arranged to initiate fracturing and propagation of skin tearing. By preconditioning the tissue for autotomy, the fracture lattice dampens the damage-induced inflammatory response but also upregulates a pro-regenerative gene signature, accelerating skin appendage regeneration. Lastly, we discovered the key role of spiny hairs in fracture lattice formation, as inhibiting their development leads to abnormal pattern formation and changes in skin fracture mechanics. Our results present a novel example of a uniquely evolved structural adaptation in mammalian skin that links tissue patterning, autotomy and regeneration. We expect that the application of a modular compartment structure to artificial skin and other organ engineering may enhance resilience to injury and facilitate efficient regeneration.

## Main text

Shedding body parts (autotomy) is a unique adaptation that allows an organism to swiftly detach a body part to elude capture^1–3^. Due to its high biological costs^15^, autotomy must be tightly regulated: it should occur in the presence of physical threat, and the detachment must be rapid and limited to a specific body part^4,5^. Lizards are widely known for tail autotomy where the tail can release along preformed planes of weakness called fracture planes^6–9^. Although well-documented in reptiles, autotomy among mammals is rare. Intriguingly, spiny mice (*Acomys*) are one such exception, exhibiting tail and skin autotomy^10,11^. However, the remarkable ability of autotomy comes with a trade-off: *Acomys* skin is mechanically weak^10,12^.

Like lizards, *Acomys* has evolved an elegant solution. After skin tearing, this rodent exhibits exceptionally efficient and complete tissue regeneration^16,17^. In most other mammals, including humans, skin wound healing typically results in fibrotic scar tissue that is deficient in function and devoid of the original tissue architecture^18,19^. By contrast, *Acomys* demonstrates an unparalleled capacity for complete regeneration of skin and its appendages—restoring hair follicles, sebaceous glands, arrector pili muscles, vasculatures, adipose tissues, and sympathetic nerves^10,12–14^. This level of regenerative complexity is exceedingly rare among mammals and represents a striking example of epimorphic regeneration. Although significant progress has been made in characterizing the cellular and molecular mechanisms of *Acomys* skin regeneration^10,12–14,16,17,20^, the evolutionary origins of its remarkable regenerative capacity are still poorly understood.

One plausible explanation is that *Acomys* skin evolved superior regenerative capacity as a compensatory adaptation to its fragility. Indeed, in many invertebrates and non-mammalian vertebrates, the loss of body parts through autotomy is typically offset by robust tissue regeneration, enabling structural and functional recovery^21–24^. However, the mechanisms underlying skin autotomy in *Acomys* are largely unknown: How are the timing and location of skin autotomy in *Acomys* regulated? Does *Acomys* skin possess specialized structural features that support controlled tissue fracture? How does skin autotomy affect subsequent regenerative processes?

In this study, we identify a novel honeycomb lattice of highly ordered extracellular matrix (ECM) in *Acomys* skin that reveals unique fracture mechanics, specialized ultrastructure, and pro-regenerative features. We propose that this unique skin fracture lattice is evolutionarily designed not only for programmed tissue detachment but also for seamless regeneration. We suggest that the ultrastructure of the fracture lattice enables *Acomys* skin to undergo context-dependent detachment, thereby regulating the timing of skin autotomy. By tracing fracture lattice development, we highlight the central role of spiny hairs that play in this process and provide a novel link between tissue patterning and autotomy. Our findings shed light on how tissue modularity can contribute to superior regenerative capacity of *Acomys* skin, with potential implications for mechanical and biomedical engineering.

### The unique fracture properties of *Acomys* skin

*Acomys* skin is known to easily tear under mechanical stress as might occur during a predator’s attack (Fig. 1a)^16^. To experimentally mimic this scenario, we devised a “pinch load” skin fracture test by piercing the back skin with a clip and applying force upward (Fig. 1b, Extended Data Fig. 1a and Supplementary Video 1). Upon pulling, skin fracture occurred specifically at the regions adjacent to the point of applied stress, as indicated by the red arrow in Fig. 1b.

**Fig. 1:**
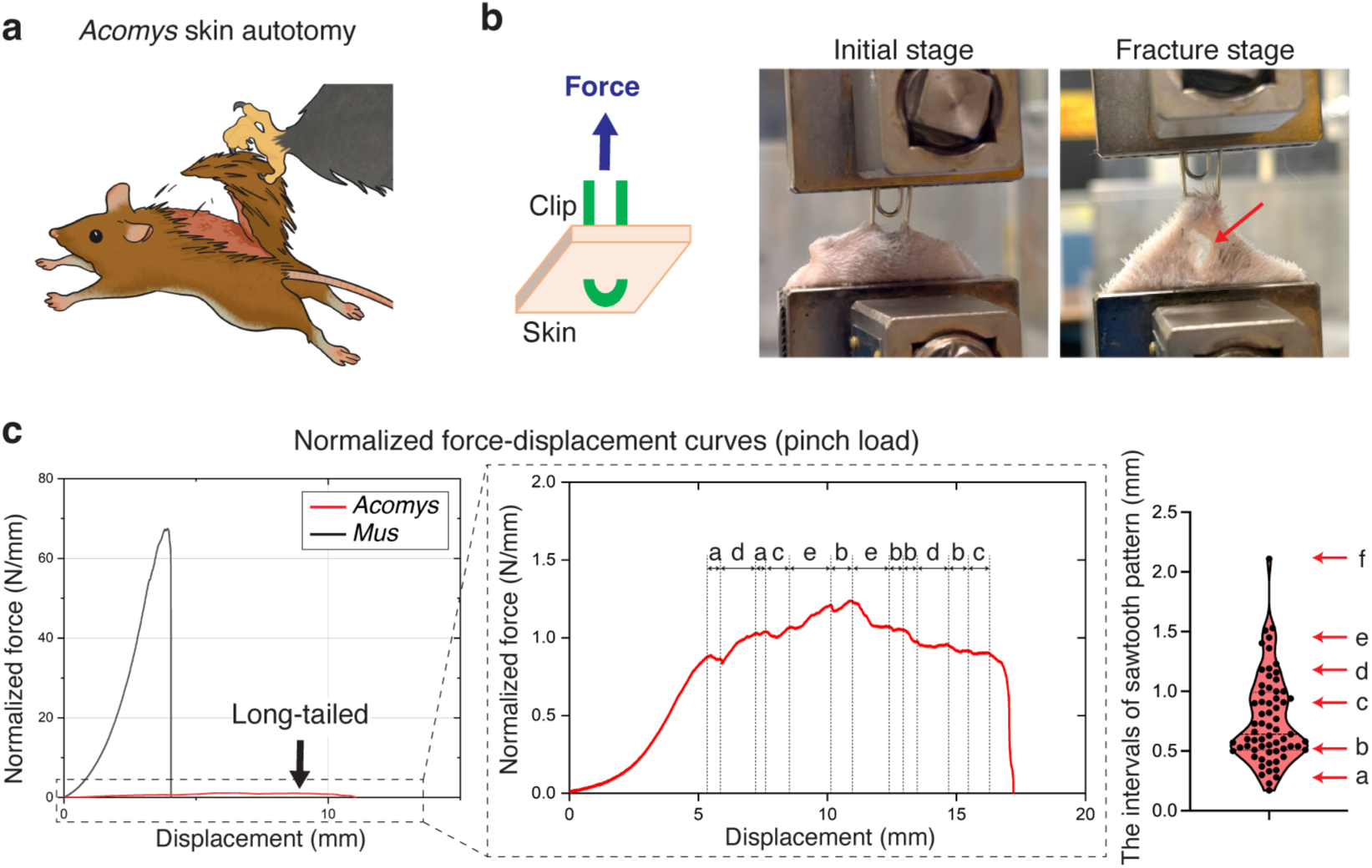
The *Acomys* skin exhibits unique mechanical properties. **a**, Schematic depicting *Acomys* skin autotomy. **b**, Illustration and photographs of a pinch load skin fracture test. Red arrow indicates skin fracture. **c**, Representative normalized force-displacement curves of *Acomys* and *Mus* skin fracture tests by a pinch load (n=5). *Acomys* data is presented in the magnified graph which exhibits a sawtooth pattern. Note that the intervals of the sawtooth pattern were not random but discrete.

Comparing force-displacement curves, we found that *Acomys* skin was ∼46 times weaker than *Mus* skin (Fig. 1c and Extended Data Fig. 1b), echoing previous findings^10^. Noticeably, in contrast to the smooth profile typically described by the worm-like chain model fracture of *Mus* skin^25,26^, *Acomys* skin showed a pronounced sawtooth-patterned fracture characterized by periodic force peaks (Fig. 1c and Extended Data Fig. 1c). This sawtooth-patterned fracture in *Acomys* skin implied that the tissue could tear gradually under a nearly constant force, akin to the action of “unzipping”. Upon further analysis, we realized that the displacement intervals observed in the sawtooth pattern were not random but discrete (Fig. 1c). The periodic fracture pattern implied the presence of a regularly repeated microstructure in *Acomys* skin.

### The regularly patterned structure in *Acomys* skin

Building on our fracture tests, we carefully characterized *Acomys* skin in multiple orientations and in doing so discovered a previously unknown repeating pattern spanning the entire back skin (Extended Data Fig. 2a). To better characterize the pattern, we performed hematoxylin and eosin (H&E) staining analyses with horizontal and vertical sections of *Acomys* and *Mus* skin (Fig. 2a). While *Mus* skin exhibited typical dermal architecture devoid of a noticeable pattern (Fig. 2b), *Acomys* skin showed a clearly demarcated pattern of repeating hexagons (short axis: 1.1 ± 0.15 mm, long axis: 1.4 ± 0.14 mm) (Fig. 2c and Extended Data Fig. 2b). Each unit was composed of 50∼100 μm thick fibrous boundaries that surrounded lipid in the lower dermis or units of three spiny hair follicles in the upper dermis (Fig. 2c and Extended Data Fig. 2b). A vertical section and whole-mount three-dimensional imaging revealed that each unit of the pattern spanned the full skin thickness and was tilted posteriorly to align with hair follicles (Fig. 2d,e and Supplementary Video 2). Immunostaining demonstrated that each unit of the pattern consisted of a collagen boundary (labelled by CNA35-EGFP, a pan-collagen probe) and three central spiny hair follicles (KRT14^+^) with adipocytes (PLIN1^+^) filling out the remaining space (Fig. 2f).

**Fig. 2:**
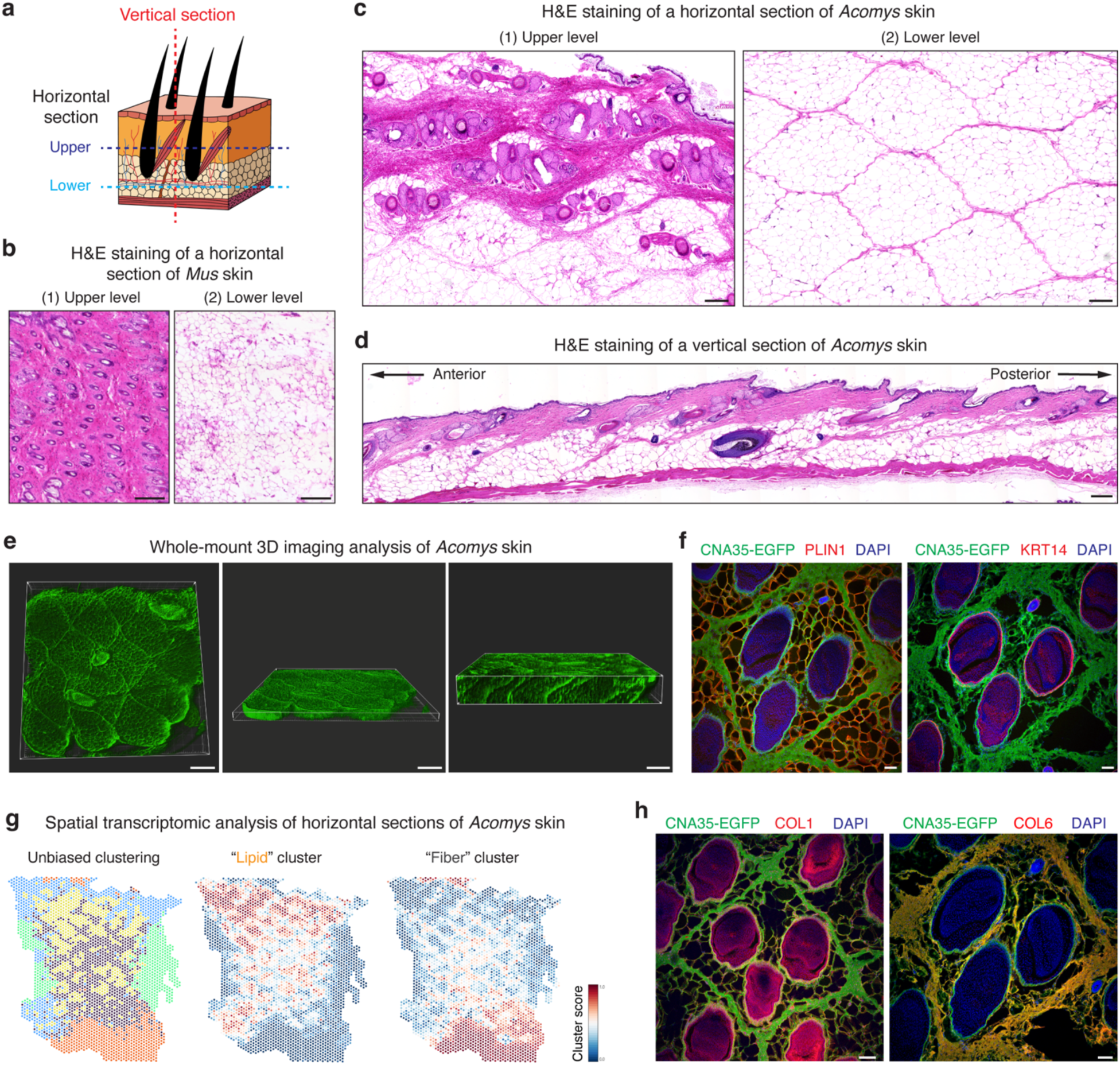
A honeycomb lattice in *Acomys* skin. **a**, Schematic depicting the orientation of the skin tissue section. **b**, H&E images showing horizontal sections of the upper and lower levels of *Mus* skin. Scale bars, 200 μm. **c**, H&E images showing horizontal sections of upper and lower levels of *Acomys* skin. Scale bars, 200 μm. **d**, H&E image showing a vertical section of *Acomys* skin. Note that each unit of the pattern spans full skin thickness and is tilted posteriorly in line with hair follicles. Scale bars, 200 μm. **e**, Representative images of whole-mount 3D visualization of *Acomys* skin. The overall structure of the skin is labeled by CNA35-EGFP, an EGFP-tagged pan-collagen binding protein. Scale bars, 500 μm. **f**, Immunofluorescence staining of horizontal sections of *Acomys* skin showing that each unit of the pattern consists of a collagen boundary (CNA35-EGFP^+^), central spiny hair follicles (KRT14^+^) and adipocytes (PLIN1^+^) padding in between. Scale bars, 200 μm. **g**, Unbiased spatial transcriptomic analysis revealed fiber and lipid clusters in *Acomys* skin. **h**, COL6, not COL1, is a major component of the collagen boundary in *Acomys* skin. Scale bar, 100 μm.

Supporting these results, spatial transcriptome analysis revealed a gene expression map reflecting the honeycomb pattern in *Acomys* skin (Fig. 2g and Extended Data Fig. 3a). Unbiased clustering separated “Lipid” and “Fiber” clusters (Fig. 2g), where the “Fiber” cluster was enriched with ECM-related genes (*Col6a2*, *Lum*, and *Loxl2*) while the “Lipid” cluster showed adipocyte-related gene expression (*Car3*, *Lpl*, and *Lep*) (Extended Data Fig. 3b.c).

Next, to gain insight into the fracture-prone nature of *Acomys* skin, we sought to characterize the ECM composition of the lattice boundaries. Unexpectedly, the boundary borders minimally expressed COL1 and COL3 (Fig. 2h and Extended Data Fig. 3d), which are generally the most abundant collagen members in skin tissues^27^. To determine which type of collagens were expressed in the boundary ECM, we used bulk RNA sequencing of *Acomys* skin and found that *Col6a1*, *Col6a2*, and *Col17a1* were highly expressed following *Col1a1*, *Col1a2*, and *Col3a1* (Extended Data Fig. 3e). By immunostaining, we found that COL6 was highly abundant in the boundary ECM (Fig. 2h), while COL17 was expressed in hair follicles (Extended Data Fig. 3d)^28^. The widespread expression of COL6 in *Acomys* skin is notable compared with *Mus* skin where its expression is confined to hair follicles and hypodermal adipocytes (Extended Data Fig. 3f). Together, these data reveal a honeycomb-like pattern in *Acomys* skin where each repeating unit surrounds three spiny hair follicles embedded in adipocytes and enclosed by the COL6-rich matrix.

### The *bona fide* fracture lattice of *Acomys* skin

We next asked if the repeating hexagonal architecture could assist rapid and effective tissue destruction in *Acomys* skin. To test this, we first conducted stress concentration simulation upon pinch loading in the skin with and without the pattern. We developed a finite element model mimicking *Acomys* skin structure based on material properties measured by tensile tests and atomic force microscopy (Extended Data Fig. 4a-c). When an out-of-plane force was applied to skin lacking a patterned structure, maximum stress was localized directly beneath the applied load (Fig. 3a and Supplementary Video 3). In contrast, in patterned skin, stress concentration along the collagen borders of the hexagon pattern increased approximately fivefold (Fig. 3a and Supplementary Video 3). This suggests that the soft lipid structure redirects stress to the collagen boundaries, which might facilitate crack initiation and propagation within the collagen fibers.

**Fig. 3:**
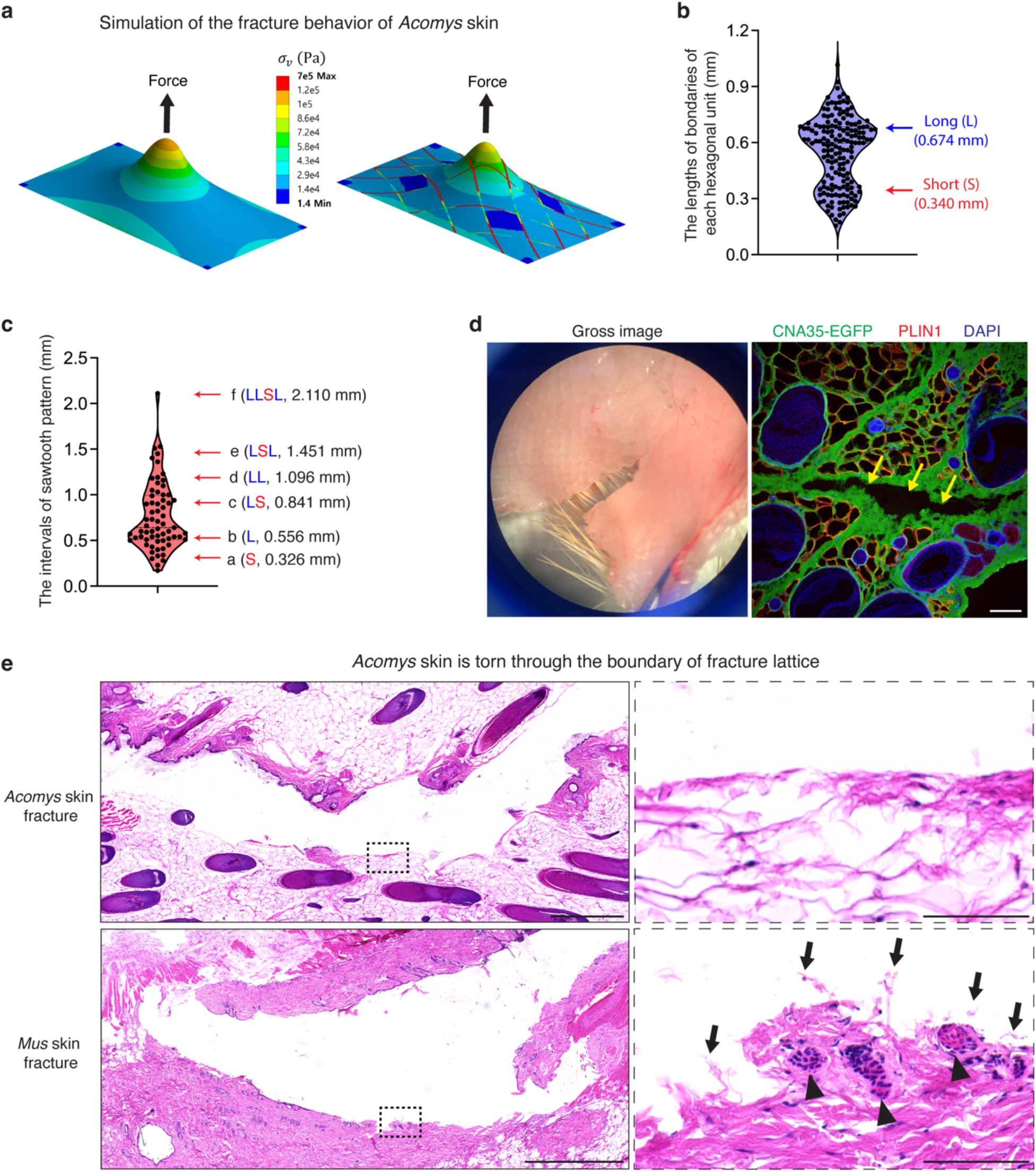
The *bona fide* fracture lattice of *Acomys* skin. **a**, The stress distribution obtained from the simulation upon pinch loading in the skin with and without the pattern suggests highest stress concentration at the collagen boundary in the presence of the pattern. **b**, A violin plot showing the lengths of edges of each hexagonal unit. **c**, A violin plot showing the intervals of sawtooth pattern. Note that the discrete pattern can be explained by the lengths of short and long boundaries of each hexagonal unit and their combinations. **d**, Photograph (left) and immunofluorescence image (right) show that *Acomys* skin tears along the collagen boundary. Scale bar, 100 μm. **e**, H&E images highlighting the torn edge of *Acomys* and *Mus* skin after the pinch load fracture test. Note that the torn edge of *Acomys* exhibits smooth collagen fibers and undamaged surrounding tissues, whereas *Mus* shows a rough edge with ruptured collagen fibers (arrows) and damaged hair follicles (arrowheads). Scale bars, 1 mm (left), 100 μm (right).

Next, we measured the dimension of each hexagonal unit and found that it consists of four long boundaries (average length, 0.674 mm) and two short boundaries (average length, 0.340 mm) (Fig. 3b and Extended Data Fig. 4d). To our surprise, the discrete intervals of the sawtooth pattern fracture analyzed in Fig. 1c could be explained by the measured lengths of long and short boundaries and their combinations (Fig. 3c). This strongly suggests that the skin autotomy takes place through the boundaries of hexagonal pattern, and the repeating pattern allows a modular opening process. Indeed, we consistently observed that *Acomys* skin tears only along the collagen boundary (Fig. 3d), resulting in a jagged-shaped edge as seen in fight skin injuries of *Acomys* (Extended Data Fig. 4e). These features suggest that the collagen boundaries along the hexagons in *Acomys* skin represent a *bona fide* fracture lattice. In contrast to a fracture plane—a continuous surface of predetermined breakage points, a fracture lattice consists of repeated structural units that facilitates tissue breakage, with break points occurring randomly depending on the strength and direction of the applied force (Extended Data Fig. 4f). In sum, unlike lizard tail autotomy, which involves a vertical fracture plane perpendicular to the tail’s axis, *Acomys* skin autotomy appears to involve a ‘fracture lattice’ oriented parallel to the skin surface that can be unzipped by an applied force.

We further analyzed the tissue architecture of the torn wound edge under high magnification. We found that the torn edges of *Acomys* skin were smooth with only a few severed collagen fibers, while *Mus* skin showed a rough edge with numerous exposed ends of severed collagen fibers (Fig. 3e), implying that collagen bundles at the fracture lattice are specifically organized for programmed tearing. Surprisingly, neighboring hair follicles and adipose tissues of *Acomys* remained structurally intact, unlike *Mus* (Fig. 3d,e), suggesting a protective role of the fracture lattice during skin autotomy. Together with the fracture simulation and histology analysis, we revealed that the collagen boundary of *Acomys* skin pattern functions as a fracture lattice by assisting crack propagation and minimizing damage to the neighboring tissue.

### Ultrastructure of the *Acomys* skin fracture lattice governs context-dependent tissue destruction

Several features of *Acomys* skin fracture in computer simulation, skin fracture tests and histological analyses hint at the hidden complexity within the fracture lattice. First, *Acomys* skin exhibited unexpected fractures in sites other than the initial crack (Extended Data Fig. 5a), suggesting that crack initiation is remarkably easy. Second, our finite element model simulation predicts collagen junctions as primary sites for stress concentration and crack initiation (Extended Data Fig. 5b and Supplementary Video 4), implying a specific structural feature underlying fracture initiation. Third, in the fracture surface of *Acomys* skin, ruptured collagen fibers were hardly seen, and the fracture seemed to take place along the grain, suggesting that collagen fibers are aligned in the same direction facilitating fracture propagation (Fig. 3e).

Based on these findings, we sought to characterize ultrastructural features of the fracture lattice. First, using structure-based super-resolution microscopy, we found two different arrangements of collagen bundles in the fracture lattice of *Acomys* skin (Extended Data Fig. 5c). Of note, the boundary ECM showed a unidirectional arrangement of fiber bundles analogous to a woven rope, while the junction ECM showed tangled fibers with few adipocytes embedded in between (Extended Data Fig. 5c,d). To further analyze the architecture in higher resolution, we visualized *Mus* and *Acomys* skin with scanning electron microscopy (SEM). Whereas *Mus* exhibited highly packed and randomly intermingled collagen fiber bundles (Fig. 4a), *Acomys* showed distinct architecture of boundary and junction collagen (Fig. 4b,c), mirroring the findings from super-resolution microscopy. Interestingly, boundary fibers were aligned horizontally, surrounding sac-like adipose tissue, while junction fibers were in perpendicular with the skin epidermis and boundary fibers (Fig. 4b,c). Structurally, boundary fibers were less compacted, curly, and associated with a large number of binding proteins (Fig. 4d, upper panels), suggestive of the elastic and dynamic nature. On the other hand, junction fibers were dense and highly packed with few associated proteins (Fig. 4d, lower panels).

**Fig. 4:**
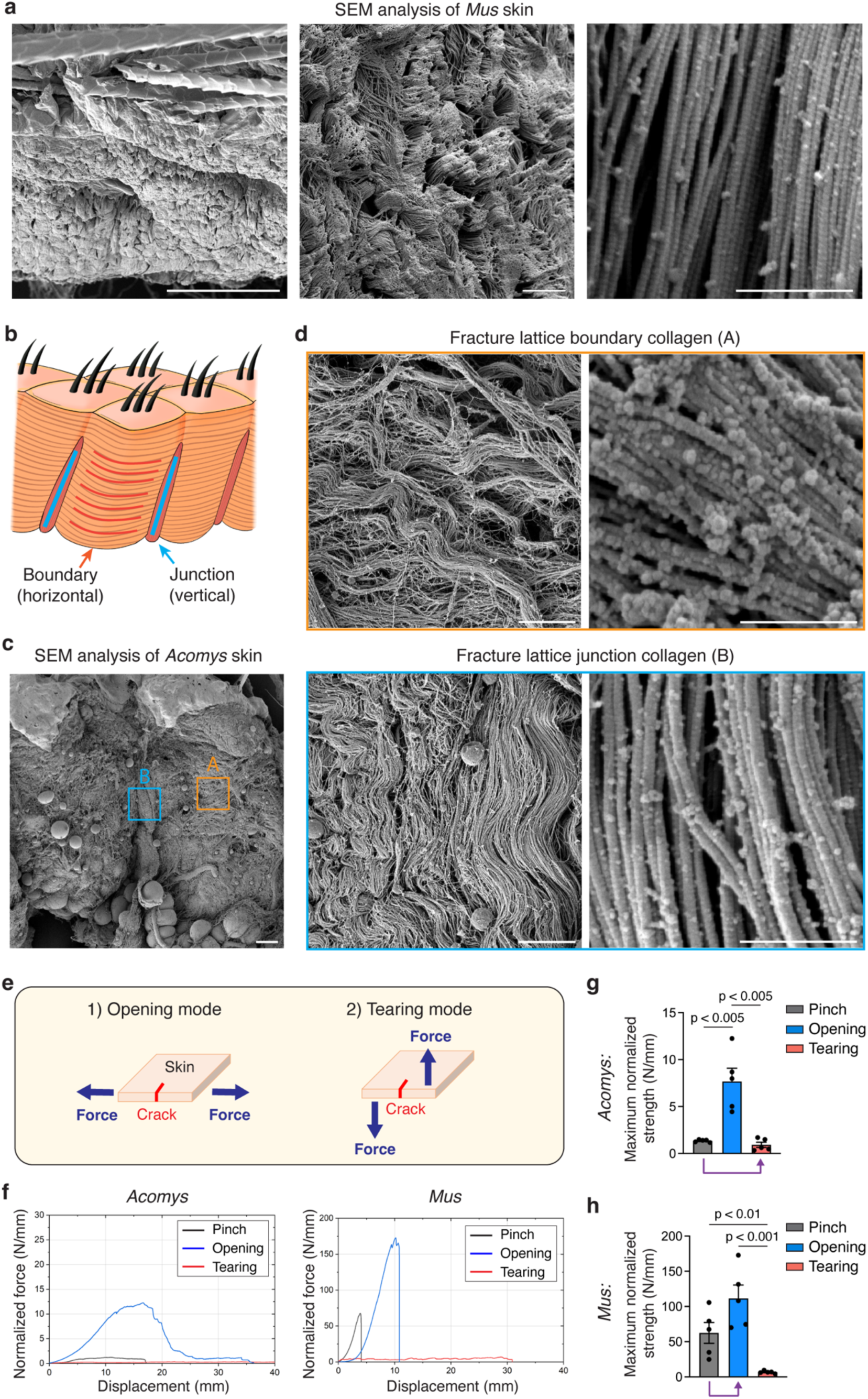
The ultrastructure of the fracture lattice of *Acomys* skin. **a**, SEM images showing densely packed and randomly oriented collagen bundles in *Mus* skin. Scale bars, 100 μm (left), 10 μm (middle), 1 μm (right). **b**, Schematic depicting the perpendicular arrangement of boundary and junction collagens in *Acomys* skin. **c**, SEM image showing the ultrastructure of *Acomys* skin. Scale bar, 100 μm. **d**, SEM images showing horizontal boundary collagen fibers (upper) and vertical junction collagen fibers (lower). Note that boundary collagen fibers are less compacted, curly, and associated with a large number of binding proteins while junction collagen fibers are dense and highly packed with few associated proteins. Scale bars, 10 μm (left), 1 μm (right). **e**, Illustrations showing skin fracture tests by the opening and tearing modes. **f**, Representative normalized force-displacement curves of *Acomys* and *Mus* skin tensile tests by the opening (n=5) and tearing (n=5) modes. **g**,**h**, Graphs comparing maximum normalized strength of three tensile tests (pinch loading, opening mode and tearing mode) in *Acomys* (**g**) and *Mus* (**h**) skin. Purple arrows indicate that pinch loading induces a tearing mode fracture in *Acomys* skin, but it resembles an opening mode fracture in *Mus* skin. For **g** and **h**, statistical analysis was performed using unpaired two-tailed Student’s t-tests. Data are mean ± s.e.m.

These microscopic boundary and junction collagen structures in *Acomys* skin point to context-dependent tissue destruction. *Acomys* skin can effectively withstand in-plane stress (horizontally), which commonly occurs in daily activities, as the stretchable horizontal boundary collagen bundles allow for effective stress dissipation (Extended Data Fig. 5e). Conversely, when subjected to out-of-plane stress (perpendicularly), as might occur during a predator’s attack, *Acomys* skin can be easily torn due to the lack of interconnections of vertical collagen bundles (Extended Data Fig. 5e). Moreover, the perpendicular arrangement of boundary and junction fibers suggests their weak adhesion, facilitating crack initiation at the junction site under out-of-plane stress (Fig. 4b,c and Extended Data Fig. 5e). This unique arrangement of collagen bundles in fracture lattice might govern context-dependent tissue tearing in *Acomys* skin.

To test this hypothesis, we set up skin fracture tests simulating two different stress conditions: (1) an opening mode, in which force is applied parallel to the tissue plane (in-plane stress) and (2) a tearing mode, in which force is applied perpendicular to the tissue plane (out-of-plane stress) (Fig. 4e, Extended Data Fig. 1a, and Supplementary Video 5 and 6). To assess the forces required for crack propagation in each mode, we developed a computer simulation model for each skin fracture mode (Supplementary Text). The simulation results revealed that crack propagation in the opening mode required greater forces compared to the tearing mode (Extended Data Fig. 5f and Supplementary Video 7). In line with this, the tensile strength in the opening mode exceeded that in the tearing mode for both species (Fig. 4f, blue curves vs red curves, and Extended Data Fig. 1b). However, we unexpectedly observed species-specific differences in how pinch load (a mixture of in-plane and out-of-plane stress) fractures relate to these two fracture modes. In *Acomys* skin, the fracture responses under pinch loading closely resembled those observed in the tearing mode (Fig. 4f,g, purple arrow), while they aligned more closely with those of the opening mode in *Mus* skin (Fig. 4f,h, purple arrow). This divergence indicates that, under pinch loading stress, *Acomys* skin undergoes immediate transition to a tearing fracture mode, while *Mus* skin primarily follows an opening fracture mode. In contrast, under in-plane stress, *Acomys* skin exhibits relatively higher resistance, due to its inherent stretchability (Fig. 4f and Supplementary Video 5). This observation supports our hypothesis that tissue destruction in *Acomys* skin is context-dependent, allowing for rapid detachment when subjected to out-of-plane stress, but resisting tissue breakage under in-plane stress. Together, our results unravel a unique evolutionary structural adaptation for conditional skin autotomy in *Acomys*, highlighting the role of fracture lattice in regulating the timing of skin autotomy.

### The regenerative functions of the fracture lattice of *Acomys* skin

Based on our evidence that the observed fracture lattice architecture promotes tearing while minimizing tissue damage (Fig. 3e), we asked if it might also accelerate regenerative healing. To test this, we performed a full thickness skin wound healing assay comparing ripped (intergranular fracture) and cut (transgranular fracture) injuries. Overall wound closure rates were comparable between the two groups (Fig 5a), but we noticed that the regenerated skin area was significantly smaller in the ripped versus cut injury (Fig. 5b). Furthermore, ripped injuries resulted in more regenerated hair follicles with increased thickness after wound healing (Fig. 5c), suggesting that the fracture lattice provides aids in tissue regeneration.

**Fig. 5:**
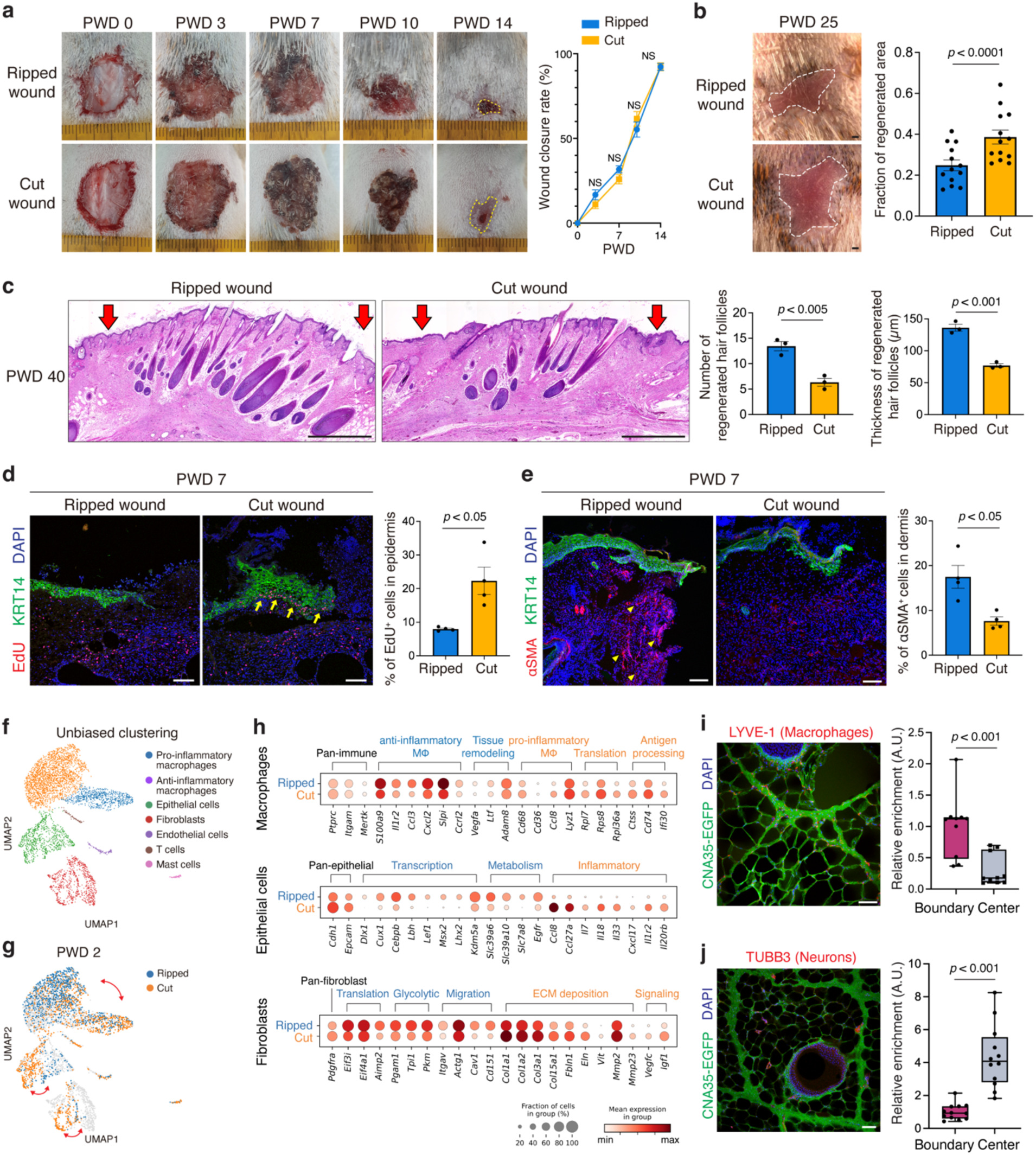
The regenerative functions of the fracture lattice of *Acomys* skin. **a**, Representative photographs and a quantification graph (n=13) show that wound closure rate is comparable between ripped (intergranular damage) and cut (transgranular damage) wounds. **b**, Photographs and a quantification graph (n=13) showing smaller regenerated area in the ripped wound at PWD 25. Scale bars, 1 mm. **c**, H&E images and quantification graphs (n=3) showing increased number and thickness of regenerated hair follicles in the ripped wound at PWD 40. Scale bars, 1 mm. **d**, Immunofluorescence images and a quantification graph (n=4) showing EdU+ proliferating cells in the wound edge regions at PWD 7. Scale bars, 100 μm. **e**, Immunofluorescence images and a quantification graph (n=4) showing αSMA+ myofibroblasts in the wound edge regions at PWD 7. Scale bars, 100 μm. **f**, UMAP projections and clustering analysis of single cell transcriptomes across wound types and days post wounding. **g**, PWD 2 shows the most divergent changes between wound types. Red arrows indicate the differences between ripped and cut injury samples. **h,** Dot plots showing differential gene expressions between ripped and cut wounds at PWD 2. The color of dots represents the mean level of gene expression, and the size of dots stands for the fraction of cells expressing each gene. **i**, Immunofluorescence image and a quantification graph showing that LYVE-1+ macrophages are enriched in the collagen boundary of the fracture lattice in *Acomys* skin. Scale bars, 100μm. **j**, Immunofluorescence image and a quantification graph showing that TUBB3+ neurons are enriched in the center regions of the fracture lattice in *Acomys* skin. Scale bars, 100μm. For **a**, **b**, **c**, **d**, **e**, **i**, and **j**, statistical analysis was performed using unpaired two-tailed Student’s t-tests. Data are mean ± s.e.m.

To investigate the underlying mechanism, we tested the extent to which dermal contraction and epithelial proliferation, two major skin wound healing processes, contributed to skin repair^29,30^. While cut injuries resulted in a higher rate of epithelial cell proliferation (Fig. 5d), ripped injuries showed an increased number of myofibroblasts (Fig. 5e), indicative of strong dermal contractility. Moreover, our computer simulation revealed a stronger contractile force concentrated at the boundary of the ripped injury (Extended Data Fig. 6a-c), suggesting that the fracture lattice structure is intrinsically designed to respond differently to the two types of skin wounds. Based on previous literature showing the relationship between tissue stiffness and wound-induced hair follicle neogenesis in *Acomys* skin^12^, we posited that the change in tissue stiffness due to dermal contraction is important for the enhanced hair follicle neogenesis in ripped injuries. To test this, we applied a splint to the back skin prior to wounding to prevent tissue contraction (Extended Data Fig. 6d). Consequently, the differences between ripped and cut injuries diminished, and the quantity and quality of newly formed hair follicles also declined (Extended Data Fig. 6e,f). This supports our hypothesis that ripped injury induces enhanced regenerative wound healing by promoting dermal contraction.

Finally, to delve deeper into molecular mechanisms, we examined gene expression differences in the acute wound healing niche at single-cell resolution. At 6 hours, 2 days, and 6 days post wounding, single cells from the wound edges of ripped and cut injuries were isolated and analyzed. Unbiased clustering analysis revealed eight distinct cell populations including pro-inflammatory macrophages, anti-inflammatory macrophages, epithelial cells, fibroblasts, endothelial cells, T cells, and mast cells (Fig. 5f and Extended Data Fig. 7a,b). Analyzing cell composition across timepoints and wound types revealed post-wound day (PWD) 2 as the most divergent timepoint, particularly showing significant differences in the percentages of pro-inflammatory and anti-inflammatory macrophages. (Fig. 5g and Extended Data Fig. 7b,c).

Differential gene expression analysis revealed distinct responses to the types of injury. In the ripped injury, there was an upregulation of genes associated with anti-inflammatory macrophages (*S100a9*, *Cxcl2*, and *Slpi*) and tissue remodeling (*Vegfa* and *Ltf*) (Fig. 5h and Extended Data Fig. 7d). Conversely, the cut injury displayed an increase in genes related to pro-inflammatory macrophages (*Cd68*, *Ccl8*, and *Lyz1*) and antigen processing (*Ctss*, *Cd74*, and *Ifi30*) (Fig. 5h and Extended Data Fig. 7d). In epithelial cells, while inflammatory response-related genes (*Ccl8*, *Il7*, *Il18*, and *Il33*) were amplified in the cut injury, early induction of key hair follicle-forming transcription factors (*Lef1*, *Cux1*, *Lhx2*, and *Msx2*) was observed in the ripped injury (Fig. 5h and Extended Data Fig. 7e). Lastly, in fibroblasts, gene expressions associated with glycolysis and migration (*Pgam1*, *Itgav*, and *Cd151*) were upregulated in the ripped injury, whereas expressions related to ECM deposition (*Col1a1*, *Col1a2*, and *Col3a1*) were elevated in the cut injury (Fig. 5h). Indeed, cut injuries at PWD 18 showed denser collagen fibers, confirming the excessive collagen production as a consequence of stronger tissue damage (Extended Data Fig. 7f). In summary, ripped injury showed a distinct wound response compared to cut injury, with less inflammatory and more pro-regenerative gene signatures across multiple cell types.

Then, how are the initial wound responses different between two wound types? Given that our fracture analysis supports the role of the fracture lattice as a structural framework reducing tissue damage upon skin autotomy (Fig. 3e), we further examined whether the localization of skin cells and tissues correlates with this pattern. Intriguingly, immunostaining analyses revealed that pro-regenerative macrophages and fibroblasts are localized near the boundaries of the fracture lattice (Fig. 5i and Extended Data Fig. 8a), whereas nerves and blood vessels are relatively sparse along the edges of the pattern (Fig. 5j and Extended Data Fig. 8b). These findings imply that macrophages and fibroblasts located at fracture sites are prepared to rapidly activate wound healing processes during skin autotomy, concurrently minimizing bleeding and nerve injury. Supporting this, neurofilament H (NEFH) and FM1-43 labeled, putative Piezo2^+^ neurons^31^ were enriched at the center of the pattern along with hair follicles (Extended Data Fig. 8c), pointing to a potential mechanism for pain attenuation during autotomy. Collectively, beyond facilitating tissue breakage, fracture lattice serves as a preconditioning architectural feature that minimizes tissue damage and enhances the regenerative potential of *Acomys* skin.

### Spiny hairs link tissue patterning and autotomy in *Acomys* skin

Lastly, we sought to understand how the fracture lattice develops in *Acomys* skin. We first asked whether the fracture lattice exists across the body or is limited to back skin.

Interestingly, the skin of head, neck, vibrissa, instep, and footpad did not exhibit an apparent fracture lattice pattern, while the abdominal skin displayed only faint outlines of the pattern (Fig. 6a), suggesting that fracture lattice formation is restricted to the back skin, coinciding with the presence of spiny hairs. Next, we traced the emergence of the fracture lattice during postnatal skin development. During early hair morphogenesis (postnatal day 1 and 7), no apparent pattern was observed (Extended Data Fig. 9a). However, at postnatal day 21 when a hair cycle reaches its first resting phase, the fracture lattice became discernible in a primitive form, reflecting its initial developmental phase (Fig. 6b,c and Extended Data Fig. 9a,b). Interestingly, along with hair cycle progression, the fracture lattice pattern underwent dynamic remodeling, eventually culminating in a definitive adult form by postnatal day 77 (Fig. 6b,c and Extended Data Fig. 9b). Vertical H&E sections also clearly showed that growing spiny hair follicles at postnatal day 24 disrupt the boundaries of the primitive lattice pattern and rebuild a definitive lattice pattern (Fig. 6d). Taken together, these data indicate a potential role of spiny hair follicles in orchestrating the formation and maturation of the fracture lattice in *Acomys* back skin.

**Fig. 6:**
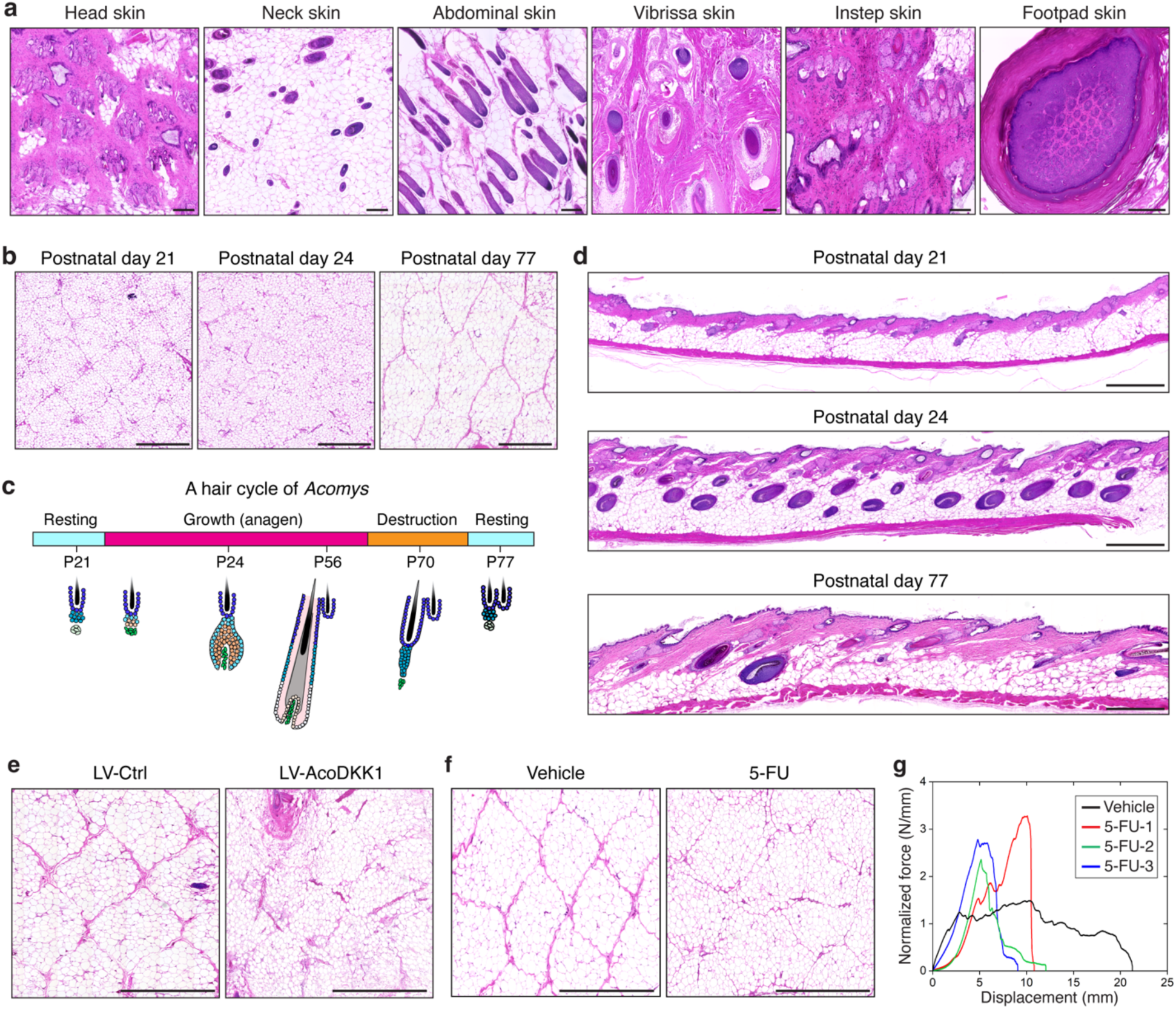
The regional specificity and temporal development of the fracture lattice in *Acomys*skin. **a**, H&E images of horizontal sections of head, neck, abdominal, vibrissa, instep, and footpad skin of *Acomys*. Scale bars, 200 μm. Note that except for a faint pattern in the abdominal skin, the fracture lattice is absent in other skin regions. **b**, H&E images of horizontal sections of *Acomys* back skin at different developmental stages. Note that at postnatal day 21, the pattern looks weak and primitive, but at postnatal day 77, the patterns become clear and definitive. Scale bars, 1 mm. **c**, Schematic depicting the hair cycle of *Acomys*. **d**, H&E images of vertical sections of *Acomys* back skin show the maturation process of the fracture lattice along with the entry of hair cycle of spiny hairs. Scale bars, 1 mm. **e**, H&E images of horizontal sections of *Acomys* back skin showing that overexpression of DKK1 disrupted the fracture lattice formation. Scale bars, 1 mm. **f**, H&E images of horizontal sections of *Acomys* back skin showing that 5-FU treatment resulted in abnormal formation of the fracture lattice. Scale bars, 1 mm. **g**, Normalized force-displacement curves of 5-FU treated and control *Acomys* skin.

To investigate the potential causal link between fracture lattice formation and spiny hair development, we employed genetic and pharmacologic strategies to inhibit spiny hair formation. First, as the WNT signaling pathway is essential for hair follicle development^32^, we overexpressed *Acomys* DKK1 in the skin epithelium using *in utero* lentiviral transduction^33^. We generated a lentiviral construct harboring expression cassettes for *Acomys Dkk1-myc-tag* and *H2b-rfp*, produced high titer lentivirus, and injected it into the amniotic sacs of *Acomys* embryos (Extended Data Fig. 9c,d)^33^. Accounting for *Acomys*’ slow embryonic development, we adjusted the timing of *in utero* injection and identified embryonic day (E) 16.5 as the optimal stage, as indicated by the robust H2B-RFP expression in the skin at E30.5 (Extended Data Fig. 9d). As previously reported^32^, DKK1 overexpression in *Acomys* epidermis efficiently inhibited hair follicle formation (Extended Data Fig. 9e). Interestingly, skin overexpressing DKK1 exhibited disrupted and incomplete fracture lattice formation, in contrast to the well-defined lattice patterning in control skin (Fig. 6e and Extended Data Fig. 9f). This result revealed a functional link between spiny hair follicle development and fracture lattice formation. Second, we topically applied 5-fluorourail (5-FU), a cell cycle inhibitor that targets rapidly proliferating hair matrix cells, leading to the destruction of hair follicles (Extended Data Fig. 9g,h). Similar to DKK1 overexpression, 5-FU treatment inhibited spiny hair formation and disrupted the fracture lattice (Fig. 6f), further confirming the critical role of spiny hair formation in fracture lattice development.

Finally, we wondered whether the disruption of fracture lattice changes the mechanical properties and fracture behavior associated with *Acomys* skin autotomy. To test this, we conducted pinch load fracture tests comparing 5-FU treated *Acomys* skin with vehicle treated controls. Interestingly, the force-displacement curves revealed two intriguing findings (Fig. 6g). First, 5-FU treated *Acomys* skin no longer exhibited the sawtooth fracture pattern but showed a worm-like chain fracture model similar to that seen in *Mus* skin yet largely decreased force value from 70N/mm down to 2-3N/mm (Fig. 6g), confirming the role of the fracture lattice in regulating skin fracture behavior. Second, 5-FU treated *Acomys* skin showed increased mechanical strength compared to control skin (Fig. 6g), functionally demonstrating the fracture lattice as a structure that facilitates skin autotomy. Together, our genetic and pharmacological manipulations revealed that spiny hair formation plays a central role in fracture lattice development, and that its disruption results in inefficient skin autotomy, highlighting an evolutionary link between tissue patterning and autotomy.

## Discussion

Autotomy, characterized by the loss of a body part, is an adaptation to evade predation and is observed across a wide variety of species, spanning from lizards to plants. The process of autotomy occurs in different ways, including external stress (*Latrodectus variolus*)^21^, muscle contraction (*Carcinus maenas*)^22^, long-lasting neuronal activity (*Aglantha digitale*)^34^, degranulation of nerve-like cells (*Melibe leonine*)^23^, and structural changes of mutable collagen fibers (*Asterias rubens*)^24^. One of the prevalent features of autotomy is the presence of a predetermined plane. For example, the tails of lizards (*Hemidactylus flavivirids*) are built in a plug-and-socket assembly^6^, and the predefined abscission zone in Bermuda buttercup (*Oxalis pes-caprae*) consists of small cells^35^. However, such examples of autotomy and the presence of fracture planes are rarely reported in mammals.

Spiny mice (*Acomys*) are a notable exception among mammals, exhibiting both skin autotomy and complete regeneration; however, the presence of defined fracture planes has not been reported^10^. In this study, we discovered a novel structural feature in *Acomys* skin—a three-dimensional pattern of repeating hexagonal units that spans the entire dorsal skin. Histological analyses, along with mechanical simulations and fracture tests, revealed that skin autotomy in *Acomys* occurs exclusively along the boundaries of the honeycomb-like pattern. We, for the first time, introduce the term “fracture lattice” in this paper to describe this modular structural units that facilitates skin autotomy. Importantly, the unique structural organization of the fracture lattice allows it to stretch and withstand horizontally applied force but easily fracture along the perpendicular direction, enabling *Acomys* to detach its skin upon unexpected perpendicular force such as a predator’s attack. This finding is both unexpected and novel. This suggests that like other invertebrates and non-mammalian vertebrates, *Acomys* has independently evolved a specialized tissue structure for skin autotomy, representing a striking example of convergent evolution. This highlights how strong selective pressure for predator can repeatedly drive the evolution of autotomy mechanisms across diverse lineages, even in mammals.

What makes our discovery even more intriguing is the modular organization of the fracture lattice in *Acomys* skin. In contrast to a single continuous plane, fracture lattice is a three-dimensional structure composed of hexagonal units with boundaries for fracture propagation.

Due to this unique modular organization, fracture can initiate in any of the unit depending on the strength and direction of the applied force. This allows *Acomys* skin to autotomize easily upon assault from any direction and sacrifice only a minimal amount of skin. Then, how did these two distinct structures (plane vs lattice) evolve independently? Fracture planes are typically found in tails and limbs, which are one-dimensional linear structures with clear segmentation; in contrast, skin is a two-dimensional planar tissue lacking such segmentation. Thus, for efficient skin autotomy, a modular design of fracture points would be more advantageous than a single plane of breakage. Our study provides a compelling example of convergent evolution occurring in distinct biological context, leading to the emergence of structurally different but functionally analogous adaptations.

Fracture lattice may provide a link between the swift autotomy of *Acomys* skin and its complete regeneration following autotomy, including all the skin appendages. We hypothesized that the fracture lattice reduces tissue damage and accelerates healing by keeping important tissue parts away from break points and placing pro-regenerative cells in proximity. Indeed, our immunostaining data revealed that hair follicles, adipocytes, nerves, and blood vessels are located within the interior of the fracture units, allowing them to remain intact following autotomy. In contrast, pro-regenerative macrophages and fibroblasts are positioned near the boundaries of the fracture lattice, where breakage occurs. We functionally tested this hypothesis by comparing ripped injury (intergranular fracture) and cut injury (transgranular fracture) and found that that ripped injury led to enhanced regeneration, characterized by higher number of regenerated hair follicles with increased thickness. Single cell transcriptomic analyses revealed that ripped injury is associated with an attenuated pro-inflammatory response, early induction of key hair follicle-inducing factors, and elevated metabolism and protein translation. These findings suggest that, by preconditioning the tissue for autotomy, the fracture lattice plays a key role in the exceptional regenerative ability of *Acomys* skin. This provides compelling and novel insights into how tissue architecture contributes to regenerative processes, with potential implications for future tissue engineering.

In line with its superior regenerative capacity, it is noteworthy that the predominant expression of collagen VI in *Acomys* skin aligns the tensile fragility across the fracture lattice and points to its functional role in skin regeneration. In contrast to the collagen I and III fibers that are major matrix constituents of *Mus* skin, our results from bulk RNA sequencing, immunostaining and SEM analyses suggest that the fracture lattice of *Acomys* skin consists mainly of collagen VI, which forms a branched network of beaded microfibrils. Collagen VI is a heterotrimeric assembly of three alpha chains (α1, α2, and α3)^36^. Particularly, Col6α3 chain has a high protein binding capacity with nine Von Willebrand factor type A domains, one fibronectin type III domain, and one Ku domain^36^, as seen in the highly decorated collagen bundles of *Acomys* skin (Fig. 4d). Given its high protein binding capacity, collagen VI is expected to promote the wound healing process in the following ways. First, upon tissue injury, collagen VI can sequester damage associated molecule patterns (DAMPs) via its protein binding domains to minimize tissue damage. The sequestering role of collagen VI has been reported to effectively reduce amyloid β’s toxicity in neuronal cells^37^. Second, collagen VI can act as a sponge for regenerative growth factors including PDGFs and TGFs^38,39^. Thus, upon injury, these stored molecules are readily available for regenerative wound healing. Third, collagen VI can provide a regenerative ECM framework for epithelial cells, fibroblasts and immune cells to activate their proliferation, migration and secretion of growth factors to enhance tissue regeneration^40^. In the future, the proposed pro-regenerative role of collagen VI in *Acomys* skin needs to be investigated.

A notable feature of *Acomys* (spiny mice) is the presence of thick and long spiny hairs that cover the entire dorsal skin. However, intriguingly, we observed that these hairs are shed quite easily, raising questions about their effectiveness as a physical barrier. Then, what is the true function of spiny hairs in *Acomys*, and how did they evolve? Skin patterning is one of the most well-known examples of Turing’s reaction-diffusion theory of tissue morphogenesis^41,42^. During embryonic skin development, the dynamic interaction between a WNT activator and its inhibitor spontaneously generates a hexagonal pattern of hair follicle spacing^43^, reminiscent of the honeycomb-like fracture lattice observed in *Acomys* skin. Notably, each hexagonal unit of the fracture lattice always surrounds three spiny hairs. Moreover, in regions lacking spiny hairs such as the head, neck, ears, and tail, the fracture lattice pattern is absent, suggesting a developmental parallel between spiny hair formation and fracture lattice development. Indeed, our genetic and pharmacological manipulations revealed that inhibiting spiny hair formation results in an incomplete formation of the fracture lattice, highlighting the central role of spiny hair formation in the establishment of fracture lattice. Although our results offer novel developmental insights into the evolutionary link between tissue patterning and autotomy, several questions on evolutionary functions remain. For example, it remains unclear whether hair enlargement, a key trait of spiny hair, is necessary for the development of the fracture lattice in *Acomys*. In fact, our histological analysis on human scalp skin (Extended Data Fig. 10) and previous studies on hair follicles in hedgehog and porcupine^44,45^ indicate that hair thickening is not sufficient to drive fracture lattice formation. This implies that additional *Acomys*-specific evolutionary innovations have occurred. Further comprehensive studies on regenerative species and unique skin appendage may answer whether linkage of tissue patterning and regeneration is solely unique feature of *Acomys* or overlooked prevalent coevolution.

In *Mus* skin wounds, tissue injury activates the hair cycle which propagates from the wound edge to the surroundings in a wave-like process (Extended Data Fig. 11). This is an injury-induced adaptation to compensate for the weakened skin barrier, in expense of high energy consumption^46^. However, this cascade of wound-induced hair cycle activation is hardly seen in *Acomys* skin (Extended Data Fig. 11). Since the threshold of hair cycle activation in *Acomys* seems comparable to *Mus*^20^, we speculate that the fracture lattice of *Acomys* skin may function as a biophysical barrier that insulates damage-associated signals. This kind of damage insulating system is found in the compartmentalized structure of spider appendages and injury-induced lignin accumulation in plants^47,48^. Analogous to the watertight subdivision of a ship’s hull in engineering, *Acomys* skin serves as an example of compartmentalization adapted for mammalian injury resistance. In the future, we expect that application of modular compartment structure to artificial skin and other organ engineering may enhance resilience to injury and facilitate efficient replacement-based repair.

In sum, this study reveals a fracture lattice in *Acomys* skin and highlights its unique characteristics including the collagen VI-rich composition, modular organization for efficient autotomy, unique ultrastructure for context-dependent skin tearing, spiny hair-driven tissue patterning, and role in regeneration. These findings provide invaluable insights into the high regenerative capacity of *Acomys* and its evolutionary basis and suggest that the application of modularity could benefit future tissue engineering research.

## Method

### Mice and procedures

Mice maintenance and experimental procedures were approved by the Institutional Animal Care and Use Committee (IACUC) of KAIST (KA2024-109-v1) and the United Kingdom Home Office. To perform ripped and cut skin wounding experiments, *Acomys* aged 12 to 20 weeks were anesthetized with isoflurane. After shaving the hair on the dorsal skin, a full-thickness wound measuring 2 to 3 cm^2^ was carefully generated by forceps along the fracture lattice (ripped wounded group), while a wound of the same shape and size was made using scissors, ignoring the fracture lattice (cut wounded group). For splinting wound regions, a splint was fixed to the shaved dorsal skin using sutures, and ripped and cut wounds were applied. For the wound-induced hair cycle activation test, a full-thickness cut wound with a diameter of 1cm was made on the back skin of 8-week-old *Mus*, while a full-thickness wound with a diameter of 1.5 cm was generated on the back skin of *Acomys*. Gross images were captured using a mobile phone (Samsung). Images of regenerated dorsal skin were taken using a stereomicroscope (SMZ18; Nikon). To label proliferating cells, EdU (25 μg/g) was injected intraperitoneally 6 hours before lethal administration of CO2.

### *Acomys* and *Mus* skin fracture tests

To examine the fracture mechanics of *Acomys* and *Mus* skin, fracture tests were conducted using a universal testing machine (EZ-SX, Shimadzu, Japan) under three different loading conditions. For a pinch load test that mimics the mechanical stress experienced by the skin during a predator’s attack, two perforations were created in the skin using clips with a 10 mm gap between them, and an out-of-plane load was applied. For an opening mode fracture test, a 6-mm pre-crack was introduced, and in-plane load was applied perpendicular to the crack face. For a tearing mode fracture test, a 6-mm pre-crack was introduced, and an out-of-plane load was applied to induce shear stress along the crack front. *Acomys* skin samples measured 30 mm × 30 mm with a mean thickness of 1.02 mm, and *Mus* skin samples measured 20 mm × 20 mm with a mean thickness of 0.23 mm. For pinch load test on 5-FU treated *Acomys,* skin samples measured 20mm x 20mm and thickness was measured separately for each sample for force normalization. The displacement rate was set to 10 mm/min for all tests. The measured load was normalized by the sample thickness.

### Hematoxylin and eosin (H&E) staining

Tissue samples were dissected and fixed overnight in 4% paraformaldehyde (PFA; EMS) and subsequently embedded in paraffin. The paraffin-embedded blocks were sectioned to a thickness of 5 μm and mounted on slides. The tissue sections were then deparaffinized and rehydrated.

Following this, the sections were incubated with hematoxylin (Sigma Aldrich) and subsequently stained with eosin (Sigma Aldrich). Images were captured using Axio Scan. Z1 (Carl Zeiss Inc). The regenerated area and hair follicle thickness were measured with ImageJ software.

### Immunofluorescence and microscopy

For immunofluorescence analysis, mouse dorsal skins were fixed with 1% PFA for 1 hour at 4℃, incubated overnight with 30% sucrose (Daejung), and then embedded in optimal cutting temperature (OCT) compound (Tissue-Tek; Sakura). Frozen tissue blocks were sectioned at 15 μm or 20 μm, and mounted on slides. If post-fixation was necessary, the samples were incubated in 1% PFA for 10 min at room temperature. Tissue sections were washed three times with PBS and blocked in blocking solution (1% bovine serum albumin (LPS solution), 1% gelatin (Sigma-Aldrich), 2.5% normal donkey serum (Jackson ImmunoResearch) and 0.3% Triton X-100 (LPS solution) in PBS) for 1 hour at room temperature. Sections were incubated overnight at 4℃ with primary antibodies diluted in blocking solution. Sections were then washed five times with PBS and incubated with secondary antibodies in blocking solution for 1 hour at room temperature.

Finally, sections were washed five times with PBS and mounted with Mounting Medium (ibidi). The following antibodies and dilutions were used: anti-PLIN1 (1:200, abcam, Cat#ab3526), anti-COL1 (1:1000, abcam, Cat#ab34710), anti-COL6 (1:200, abcam, Cat#ab182744), anti-PECAM-1 (1:200, R&D systems, Cat#AF3628), anti-K14 (1:1000, Biolegend, Cat#906004), anti-COL3 (1:200, abcam, Cat#ab7778), anti-COL17 (1:200, abcam, Cat#ab184996), CNA35-EGFP (1:2000), anti-αSMA (1:500, abcam, Cat#ab5694), anti-LYVE-1 (1:200, AngioBio, Cat#11-034), anti-TUJ1 (1:200, abcam, Cat#ab18207), anti-PDGFRa (1:200, abcam, Cat#ab203491), and anti-CUX1 (1:100, Cat#11733-1-AP). Alexa Fluor-488, −594 or −647-conjugated secondary antibodies (Life Technologies, 1:1000) were used. Nuclei were stained using 4’6’-diamidino-2-phenylindole (DAPI; Sigma-Aldrich). EdU Click-iT reaction was performed according to manufacturer’s instructions (Thermo Fisher Scientific). Images were collected using a confocal microscope (ECLIPSE Ti2; Nikon) equipped with laser wavelengths of 405, 488, 561 and 640 nm, and processed using ImageJ. For high-resolution imaging, 3D-Structured Illumination Microscopy (SIM) images were captured using a Ti-2 inverted microscope (Nikon) equipped with a Ziva Light Engine (Lumencor) as the excitation light source. This setup allows for high-resolution imaging with enhanced optical sectioning capabilities. The Ziva Light Engine provides 405, 488, 561, and 640 nm lasers for excitation. The raw dataset comprised 15 individual images, acquired under five distinct illumination patterns and three different angles of illumination, which allows for effective reconstruction of high-quality 3D images. Sequential image acquisition was carried out according to the excitation channel using a high-resolution sCMOS camera (Hamamatsu Photonics K.K., ORCA-Flash4.0 sCMOS, 2048 × 2048 pixels), which offers a large field of view and excellent sensitivity.

The acquired 3D-SIM images were then reconstructed using the NIS-Elements (Nikon) software, which employs advanced computational algorithms to produce the final high-resolution 3D images with improved spatial resolution.

### Wholemount 3D imaging analysis of *Acomys* skin

For whole-mount 3D imaging analysis, *Acomys* backskins were dissected and fixed with dimethyl sulfoxide (DMSO):methanol (1:4) for overnight at −20°C. Fixed samples were then washed in methanol:PBS+0.3% Triton X-100 (PBT) and PBT for 1 hour each at room temperature. Skin samples were blocked in blocking solution (1% BSA, 1% Gelatin, 2.5% Normal donkey serum, 0.3% Triton X-100 in PBS) for overnight at room temperature. CNA35-EGFP was diluted (1:500) in blocking solution and skin samples were incubated for 3 days at 4°C with shaking. Then, skin samples were washed in blocking solution six times for 30 minutes each at room temperature. Skin samples were gradually dehydrated in 50% and 70% ethanol for 1 hour and dehydrated in 100% ethanol twice for 40 minutes each. Finally, dehydrated skin samples were stored in ethyl cinnamate at room temperature for tissue clearing. Images were acquired with an confocal microscope (Nikon) using 4× air objective with z-stacks that were 5 µm apart. Images were processed using Imaris.

For wholemount visualization of nerves in *Acomys* skin, 13-week-old adult female spiny mice were intraperitoneally injected with FM1-43FX fixable analog (ThermoFisher, F35355) and returned to the homecage for 24 hours. Animals were then transcardially perfused with 4% PFA, skin tissue was removed and flattened overnight in fixative before being returned to phosphate buffer. Skin sections were blocked (10% NGS, 0.3% Triton X-100) and incubated overnight with anti-Neurofilament heavy polypeptide antibody (1:1000, abcam, Cat#EPR20020) with 1% NGS and 0.3% Triton X-100, followed by washing and incubation with Alexa 647-conjugated secondary antibody with 1% NGS and 0.3% Triton X-100. For top-view imaging, skin sections were mounted on slides in Fluoromount with DAPI. Embedded skin was cross-sectioned with a vibratome and then mounted on slides. Image was performed with a Zeiss Axio Imager Z2 microscope.

### Spatial transcriptomics

Two skin tissue sections from *Acomys* were processed with the standard Visium Gene Expression Slide & Reagents kit as per the manufacturer’s instructions (CG0000239, Rev F). Briefly, fresh frozen tissue sections were placed directly on Visium slides (standard), imaged without H&E staining then processed as manufacture’s instruction. Visium libraries were analyzed via quantitative polymerase chain reaction (qPCR) (CFX Opus 96, Bio-Rad), and a total of 16 (standard workflow) PCR cycles were used for library amplification. Visium libraries were quantified with Qubit 1× double-stranded DNA high-sensitivity assay kit (Invitrogen, catalog no. Q33231) on a Qubit Flex Fluorometer (Invitrogen). Qualitative assessment of Visium libraries was conducted with the Agilent High Sensitivity D5000 ScreenTape (Agilent) to assess size distribution. All Visium libraries were normalized, pooled, spiked with PhiX Control v3 (Illumina) and sequenced paired-end (300 cycle, Rd1: 150, i72: 10, i5: 10, and Rd2: 150) on a Novaseq 6000 (Illumina).

### Spatial transcriptomics analysis

The Space Ranger computational pipeline from 10x Genomics was used to generate count matrices (v2.1.0). The reads were aligned to Acomys genome (GenBank: JAULSH000000000). After preprocessing, analysis of the spatial Visium RNA-seq data was performed with the Python packages SCANPY (v.1.9.6)^49^ and Squidpy (v.1.4.1)^50^. The Space Ranger output files and the corresponding histology images were merged into a single AnnData object. Quality control parameters were assessed, but the data was not filtered to maintain low complexity adipocyte-rich spots. Log-transformed data via SCANPY’s pp.log1p() function was used to calculate highly variable genes (SCANPY default setting).

### Atomic force microscopy

The mechanical properties of the prepared *Acomys* specimens were estimated in deionized (DI) water using a commercially available atomic force microscope (Cypher-ES, Asylum Research, CA, USA). Specimen topography images were collected using chemically inert conductive diamond tips (CDT-FMR, NanoWorld, Switzerland) with a resonance frequency of 105 kHz, a measured spring constant of 8.32 N m^-1^, and a tip radius of 150 nm. Nanoindentation was conducted using colloidal silicon dioxide tips (CP-qp-CONT-SiO, sQube, Germany) with a resonance frequency of 30 kHz, a measured spring constant of 0.142 N m^-1^, and a sphere diameter of 6.62 μm. Force-distance curves were collected and analyzed using IGOR Pro 6.37 software (Wavemetrics, OR, USA), where the Hertz model was applied to estimate the sample elastic moduli, with a power law of 1.5 used to account for tip geometry, and a Poisson ratio of 0.33 assumed for the samples.

### Measurement of tensile mechanical properties of *Acomys* skin

A custom-built micro-tensile tester was used to measure the tensile mechanical properties of *Acomys* skin (Extended Data Fig. 4b). Briefly, when tensile force is applied to the specimen via a piezoelectric actuator, the leaf spring undergoes deflection. The force exerted on the specimen is calculated based on the stiffness of the leaf spring and its deflection measured by a displacement sensor. Further details regarding the micro-tensile tester can be found in previous literature^51^.

The specimens with dimensions of 20mm in length and 5mm in width were prepared for tensile testing. The specimen thickness varied between 1 and 2mm, depending on samples. Each specimen was mounted onto the micro-tensile tester and securely fastened. Tensile loading was then applied, and the resulting stress-strain curve was obtained. The tensile tests were conducted at a strain rate of 0.001 s^−1^.

### Finite element analyses

#### 1) Simulation of an opening mode and a tearing mode fracture

Finite element analysis (FEA) was conducted using the commercial software ANSYS 17.2 to investigate the fracture behavior of *Acomys* skin. The skin structure was modeled to replicate the collagen-lipid composite observed in the actual tissue, where collagen fibers were represented as thin and stiff layers surrounding the compliant lipid clusters arranged in a parallelogram configuration. Material properties used in simulations are referred to Supplementary Text. In-plane and out-of-plane loading conditions were applied to simulate mechanical stress in the opening and tearing mode, respectively. In-plane tensile loading was applied laterally to induce crack initiation and propagation, and for out-of-plane loading, a vertical displacement was imposed at the center of the model with fixed boundary conditions at the boundaries (Extended Data Fig. 4a). Further details on model parameters, meshing strategies, and validation procedures are provided in Supplementary Text.

#### 2) Simulation of mechanical stress distribution around ripped wound and cut wound

A two-dimensional finite element model was constructed in COMSOL Multiphysics (v6.2) to simulate stress redistribution in a collagen-rich extracellular matrix under mechanical deformation. The model geometry consisted of a rectangular sheet (59.701 mm × 39.442 mm) tessellated into a periodic honeycomb lattice, representing the aligned collagen fiber network. Each hexagonal unit cell had a long diagonal of 0.674 mm and a short diagonal of 0.34 mm. Two materials were defined to capture the biphasic nature of skin tissue. Collagen was modeled as a linear elastic material with a Young’s modulus of 92.2 kPa, Poisson’s ratio of 0.3, and density of 1120 kg/m³. Fat tissue was assigned a Young’s modulus of 9.4 kPa, Poisson’s ratio of 0.48, and density of 900 kg/m³. To simulate intrinsic mechanical tension within the tissue, we applied a displacement boundary condition equivalent to 10% of the hexagon length to each external edge of the structure. To assess the mechanical consequences of structural discontinuities, two central void geometries were introduced: a smooth circular cut (Diameter = 15 mm), and 254 units defected variant representing rupture wound. These regions were treated as voids with no material assignment. von Mises stress distributions were computed under quasi-static, plane-stress conditions. For quantitative boundary analysis, explicitly selected edge sets were used to extract line profiles of stress magnitude along defect perimeters. Mesh independence was confirmed when maximal stress values varied less than 1% upon further refinement.

### Scanning electron microscopy (SEM)

The samples were fixed in 2.5% paraformaldehyde-glutaraldehyde mixture buffered with 0.1M phosphate (pH 7.2) for 4 hours, postfixed in 1% osmium tetroxide in the same buffer for 1 hour, dehydrated in graded ethanol, and substituted by isoamyl acetate. Then samples were dried at the critical point in CO2. Finally, the samples were sputtered with gold in a sputter coater (SC502, POLARON) and observed using the scanning electron microscope, FEI Quanta 250 FEG (FEI, USA) installed in Korea Research Institute of Bioscience and Biotechnology (KRIBB).

### Single cell cDNA synthesis and sequencing library generation

Single cell RNA sequencing libraries were generated with minor modifications to the Smart-seq2 protocol^52^. Single cells were isolated into 96 well PCR plates (Thermo Fisher Scientific) with 2 μl of lysis buffer (0.1% Triton X-100, 1 U/μl RNase Inhibitor (Enzynomics), 0.25 μM oligo-dT30VN primer) and preserved at −80℃. The plates were thawed on ice. Reverse transcription was carried out using 20 U/µl Maxima H minus transcriptase (Thermo Fisher Scientific), 1 M betaine (Sigma-Aldrich), 5 mM MgCl2 (Enzynomics), 1 μM template switching oligo, and an additional 0.8 U/μl Rnase Inhibitor. Following this, template switching reaction and PCR pre-amplification (KAPA HiFi HotStart (Roche), 18 cycles) were conducted in accordance with the established protocol. PCR products were purified using 0.6× SPRI beads, which consisted of 2% Sera-Mag Speed Beads (Cytiva), along with 1 M NaCl (Daejung), 10 mM Tris-HCl pH 8.0 (Enzynomics), 1 mM EDTA (Enzynomics), 0.01% NP40 (Sigma-Aldrich), 0.05% Sodium Azide (Sigma-Aldrich), and 22% w/v PEG 8000 (Sigma-Aldrich). The quality of cDNA libraries was evaluated using quantitative PCR with *Acomys Gapdh* primer, where the forward primer is 5′-GTCGTGGAGTCTACTGGTGTCTTCAC-3′ and the reverse primer is 5′-GTTGTCATATTTCTCGTGGTTCACACCC-3′. An amount of 50–100 pg from each cDNA library was utilized to generate the Illumina sequencing library with EZ-Tera XT DNA library preparation kits (Enzynomics). Following the final PCR amplification, the samples were combined and purified with the MinElute PCR purification kit (Qiagen). Fragments approximately 200 bp in size were selected using 0.3× and 0.6× SPRI beads and their size was confirmed by High Sensitivity DNA ScreenTape Analysis (Agilent). The pooled and size-selected libraries were sequenced paired-end reads (75 cycle, Rd1: 38, i7:8, i5:8, and Rd2: 38) on NextSeq 550 (Illumina).

### Single cell RNA sequencing analysis

Raw FASTQ were aligned to the *Acomys* genome (GenBank: JAULSH000000000) using RSEM(v.1.3.1)^53^ with STAR (v.2.7.9.a)^54^. The expression values of each gene were quantified as both raw counts and transcripts per million (TPM) using RSEM. Analysis and visualization of the data were conducted in a Python environment (v.3.10.13) built on SCANPY (v1.9.6)^49^ together with Pandas (v.2.1.4), NumPy (v.1.26.3), SciPy (v.1.11.4), scikit-learn (v.1.3.2), AnnData (v.0.10.4), matplotlib (v.3.8.2) packages. TPM and metadata matrices for 5,944 skin cells across the wound healing were loaded in SCANPY as an AnnData object. Single cell data was preprocessed to remove lowly detected genes (expressed in <1 cells) and cells with low complexity libraries (<500 genes detected). SCANPY was used to log1p normalize TPM, and highly variable genes were detected. PCA was performed on highly variable genes, with 100 components and the svd_solver using ‘arpack’ (SCANPY default setting). To construct a k-nearest neighbors graph on Euclidean distance, 12 principal components were used. Data was visualized using UMAP in SCANPY, and clustering was done using the Leiden algorithm (with a resolution setting of 0.25). Cluster resolution was chosen after iterating through resolution parameters from 0.01 to 0.5, as best capturing cell type marker distribution. Marker gene expression based on literature was used to identify clusters. SCANPY was used to visualize selected marker genes in dot plots, or as normalized TPM visualized on UMAPs.

Differential gene expression based on cluster identity was analyzed with scanpy.tl.rank_genes_groups using t-test with overestimated variance of each group (SCANPY default setting) to identify genes that varied across wound time points and conditions.

### Bulk RNA sequencing

Total RNA of *Acomys* skin was extracted by homogenized tissues lysed in Tri Reagent (Sigma) using Direct-zol RNA Miniprep kit (Zymo Research). 100ng of total RNA was used to generate Bulk RNA sequencing library following a SHERRY protocol^55^ with minor modifications.

Briefly, reverse transcription was carried out using 100 ng of RNA, 0.5 mM dNTP, 5 uM oligo-dT30VN primer, 50U Maxima H Minus Reverse Transcriptase with 1X RT Buffer (Thermo Fisher Scientific), and 10U Murine RNase Inhibitor (Enzynomics). Reverse transcription product was processed using EZ-Tera XT DNA library preparation kits (Enzynomics). Library amplified by 12 cycles PCR using KAPA HiFi DNA polymerase (Roche, KK2502), then purified with SPRI beads and confirmed by High Sensitivity DNA ScreenTape Analysis (Agilent). Library was sequenced paired-end reads (300 cycle, Rd1: 150, i7:12, i5:12, and Rd2: 150) on NovaSeq 6000 (Illumina).

### Bulk RNA sequencing data processing

Raw FASTQ were adapter and quality trimmed by Trim Galore (v.0.6.1)^56^. Trimmed FASTQ aligned to the *Acomys* genome (GenBank: JAULSH000000000) using RSEM(v.1.3.1)^53^ with STAR (v.2.7.9.a)^54^. The expression values of each gene were quantified as both raw counts and transcripts per million (TPM) using RSEM.

### Lentivirus production and in utero transduction

To generate the *Acomys Dkk1* (*AcoDkk1*) expression construct, *AcoDkk1* cDNA was amplified by PCR, and a Myc tag was added to 3’ end of *AcoDkk1* cDNA. The resulting AcoDkk1-myc construct was cloned into the LV-CMV-PGK-H2BRFP plasmid, which was subsequently used for high-titer lentivirus production and *in utero* injections.

To produce VSV-G pseudotyped lentivirus, the LV-CMV-AcoDkk1-myc-PGK-H2BRFP plasmid and helper plasmids (pMD2.G and pPAX2) were transfected into 293FT cells. After 46 hours, viral supernatant was collected and concentrated ∼2,000-fold by Amicon® Ultra-15 Centrifugal Filter and ultracentrifugation in a MLS-50 rotor (Beckman Coulter). The concentrated high-titer lentivirus was injected into the amniotic sac of E16.5 *Acomys* embryos.

### Immunoblotting

To check the expression level of AcoDKK1, unconcentrated lentiviral supernatant was mixed with 5X SDS-PAGE buffer and loaded on 12% SDS-PAGE gels. Pre-Stainded Protein Marker (LPS Solution) was used as a molecular weight marker. After gel electrophoresis, proteins were transferred to PVDF membranes (Merck). After blocking with 5% skim milk in TBST solution (1X TBS, 0.1% Tween-20) for 1 h at room temperature, the membrane was incubated with mouse anti-Myc-Tag (1:1000, Cell Signaling Cat# 2276S) at 4℃ overnight and HRP-conjugated anti-mouse IgG secondary antibody (1:1000, Jackson ImmunoResearch). The blot was visualized using SuperSignal™ West Dura Extended Duration Substrate (ThermoFisher Scientific) and a ChemiDoc MP imaging System (Bio-Rad).

### 5-Fluorouracil (5-FU) treatment

A 5% 5-FU (MedChemExpress) stock solution was prepared by dissolving the compound in DMSO (Sigma-Aldrich). This stock was subsequently diluted in commercial hand cream (Amorepacific, Korea) to formulate a 1% 5-FU cream. The cream was topically applied to the shaved dorsal skin, specifically the spiny hair region, of *Acomys* at postnatal days 23 and 26. Skin samples were collected at postnatal day 56 for H&E and skin fracture analyses.

### Human skin samples

All procedures were conducted in accordance with the ethical principles of the Declaration of Helsinki. Human scalp and non-scalp skin tissues were obtained from post-mortem adult donors (age 40–70) through the Yonsei University Medical College Surgical and Anatomy Education Center, with informed consent and IRB approval (IRB# 4-2022-0903). Within 12 hours after death, fixation was performed via femoral arterial perfusion with 10% neutral-buffered formalin through a central line. Skin samples were collected from five scalp regions (frontal, occipital, parietal, temporal, vertex) and two non-scalp regions (axillary, pubic), followed by paraffin embedding. Both horizontal and vertical sections were stained with hematoxylin and eosin (H&E) for histological analysis (Extended Data Fig. 10).

### Statistics

Statistical and graphical data analyses were conducted using Microsoft Excel, GraphPad Prism 10, and Origin 2019 software. All experiments shown were performed with a minimum of three replicates, and representative data are shown. For all measurements, three biological replicates as well as three or more technical replicates were used. To determine the significance between two groups, comparisons were carried out using an unpaired two-tailed Student’s t test.

## Acknowledgments

We thank T. Yang, B. Yoo, B. Kim, D. Kim, I. Yoon, and Y. Ee for technical support. We thank G. Y. Koh, J. Y. Kang and C. Jang for comments on the manuscript, J. Kim for providing a CNA35-EGFP pan-collagen probe, and S. Y. Lee, I. Kim, S. Roh, S. Yun, D. Lee, M. B. Aldonza, J. Lee, and J. Choi for discussions. FACS was conducted by the KAIST Stem Cell Center Flow Cytometry Core (I. Yoon, director) and all mouse work was performed in KAIST Laboratory Animal Resource Center. We also acknowledge the technical support from ANSYS Korea. This work was supported by grants from National Research Foundation of Korea (grant nos. RS-2023-00273400 to Y.C.R.; RS-2025-00520263 to G.S.; RS-2021-NR061427, RS-2024-00436657, RS-2024-00508575, and RS-2022-NR067309 to H.Y.), the Korea Health Industry Development Institute (KHIDI) (grant no. RS-2024-00406488 to J.W.O), National Institutes of Health (grant no. R01-AR070313 to A.W.S), and Suh Kyungbae Foundation (grant no. SUHF-2101005 to H.Y.)

## Author contributions

D.K., A.W.S. and H.Y. conceptualized the study. D.K., Y.C.R., J.C., E.K., H.L, G.S. and H.Y. designed the experiments and interpreted the data. D.K., Y.C.R., H.C., A.W.S., H.L., G.S. and H.Y. wrote the manuscript. J.C., H.R., J.T., C.P.L performed tensile tests and fracture computer simulation. S.J., S.R. and S.H. performed atomic force microscopy analysis. S.S. and J.W.O performed histology analysis. J.L. and J.L. performed single cell RNA-seq analysis. S.Y. and A.M.C performed whole-mount skin image analysis. S.I.C. supported animal surgery and experiments. J.A.K. performed scanning electron microscopy analysis. All authors provided input on the final manuscript.

## Competing interests

The authors declare no competing interests.

## Materials & Correspondence

Further information and requests for resources and reagents should be directed to and will be fulfilled by the corresponding author, Hanseul Yang

## Data and materials availability

Bulk RNA sequencing, single cell RNA sequencing, and spatial transcriptomic data generated from this study have been deposited in the Gene Expression Omnibus (GEO) under accession code GSE293477 (bulk RNA sequencing, https://www.ncbi.nlm.nih.gov/geo/query/acc.cgi?acc=GSE293818, accession token: wzaliswsbbclngx), GSE293817 (spatial transcriptomics, https://www.ncbi.nlm.nih.gov/geo/query/acc.cgi?acc=GSE293817, accession token: slklmiuwjtkjxmf), and GSE293818 (single cell RNA sequencing, https://www.ncbi.nlm.nih.gov/geo/query/acc.cgi?acc=GSE293477, accession token: ilqzsyimdlgjxkx).

## Code availability

Custom code for single cell RNA-seq and Visium analyses has been deposited at github (https://github.com/yanglab-kaist/daeryeok_et_al). All of the other codes are available from the corresponding author on reasonable request.

---

**Extended Data Fig. 1.**
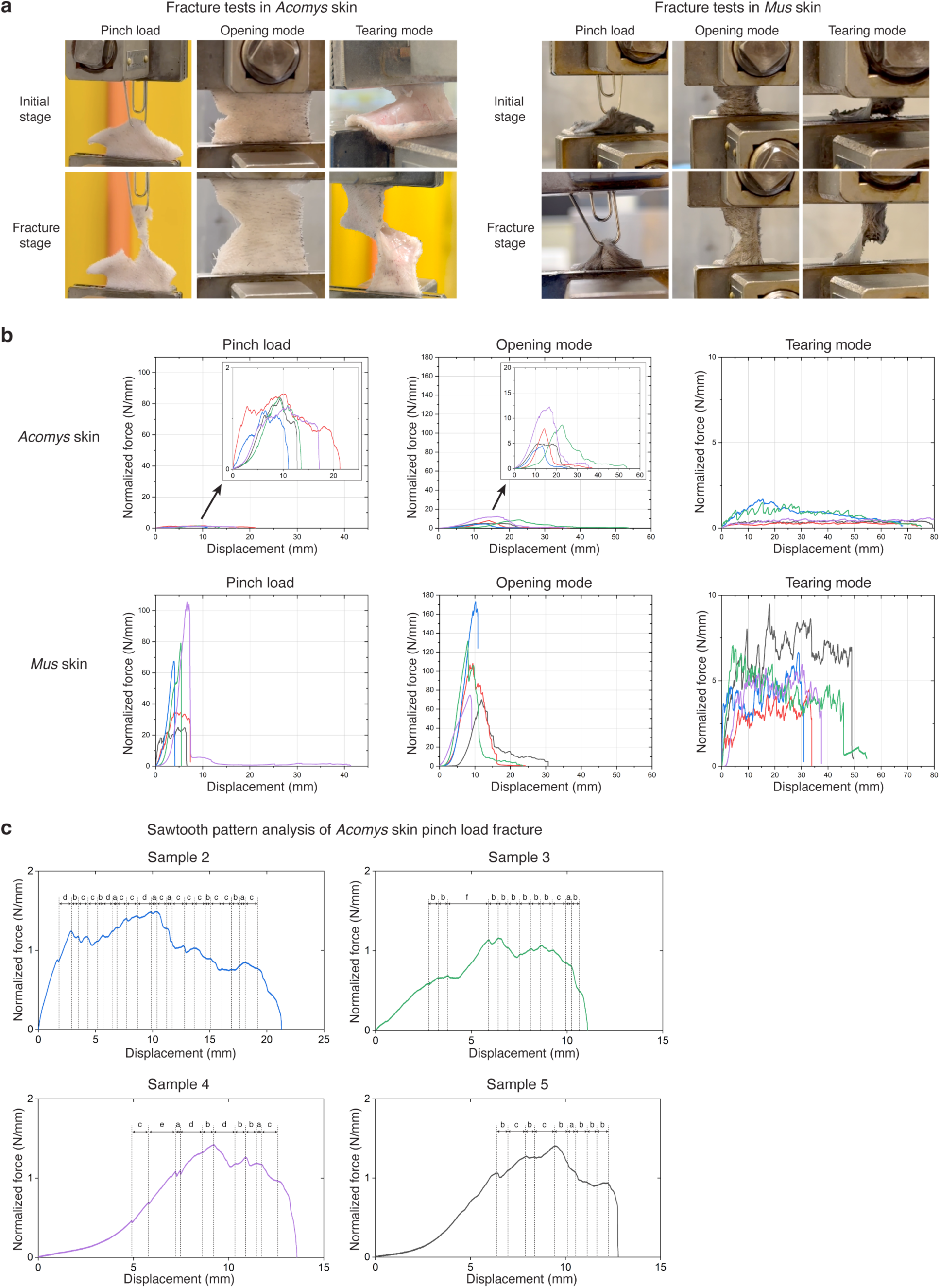
The three loadings of fracture tests reveal different mechanical properties of *Acomys* and *Mus* skin. **a**, Representative photographs showing a pinch loading, an opening mode, and a tearing mode of fracture tests at initial and fracture stages. **b**, Normalized force-displacement curves of fracture tests for *Acomys* (n=5, each) and *Mus* (n=5, each) skin. **c**, The graphs show sawtooth patterns in the normalized force-displacement curves from the pinch load fracture tests of each *Acomys* skin.

**Extended Data Fig. 2.**
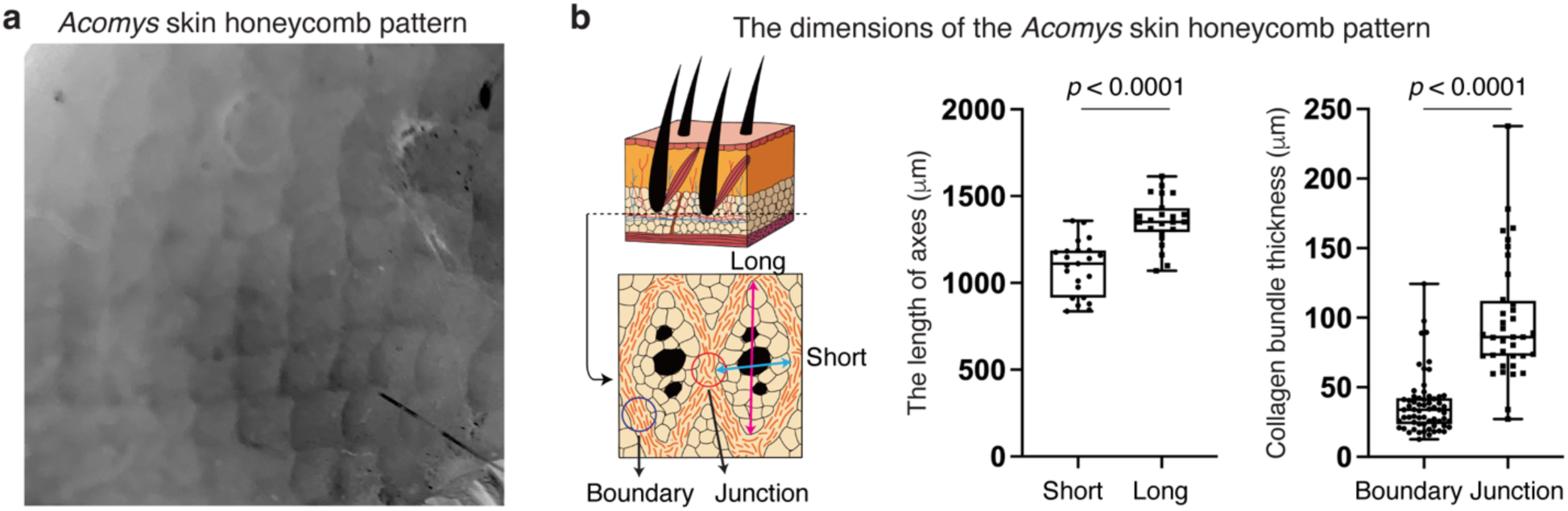
A regular and repeated pattern of *Acomys* skin. **a**, Gray-scaled photograph displaying honeycomb-like units in *Acomys* skin. **b**, Schematic depicting the boundary and junction structures of the *Acomys* skin pattern. Quantification graphs describe the dimension of the *Acomys* skin pattern. The lengths of short (n=23) and long (n=23) axes and the thickness of collagen bundles at the boundary (n=63) and junction (n=36) of the pattern were quantified. For **b**, statistical analysis was performed using unpaired two-tailed Student’s t-tests. Data are mean ± s.e.m.

**Extended Data Fig. 3.**
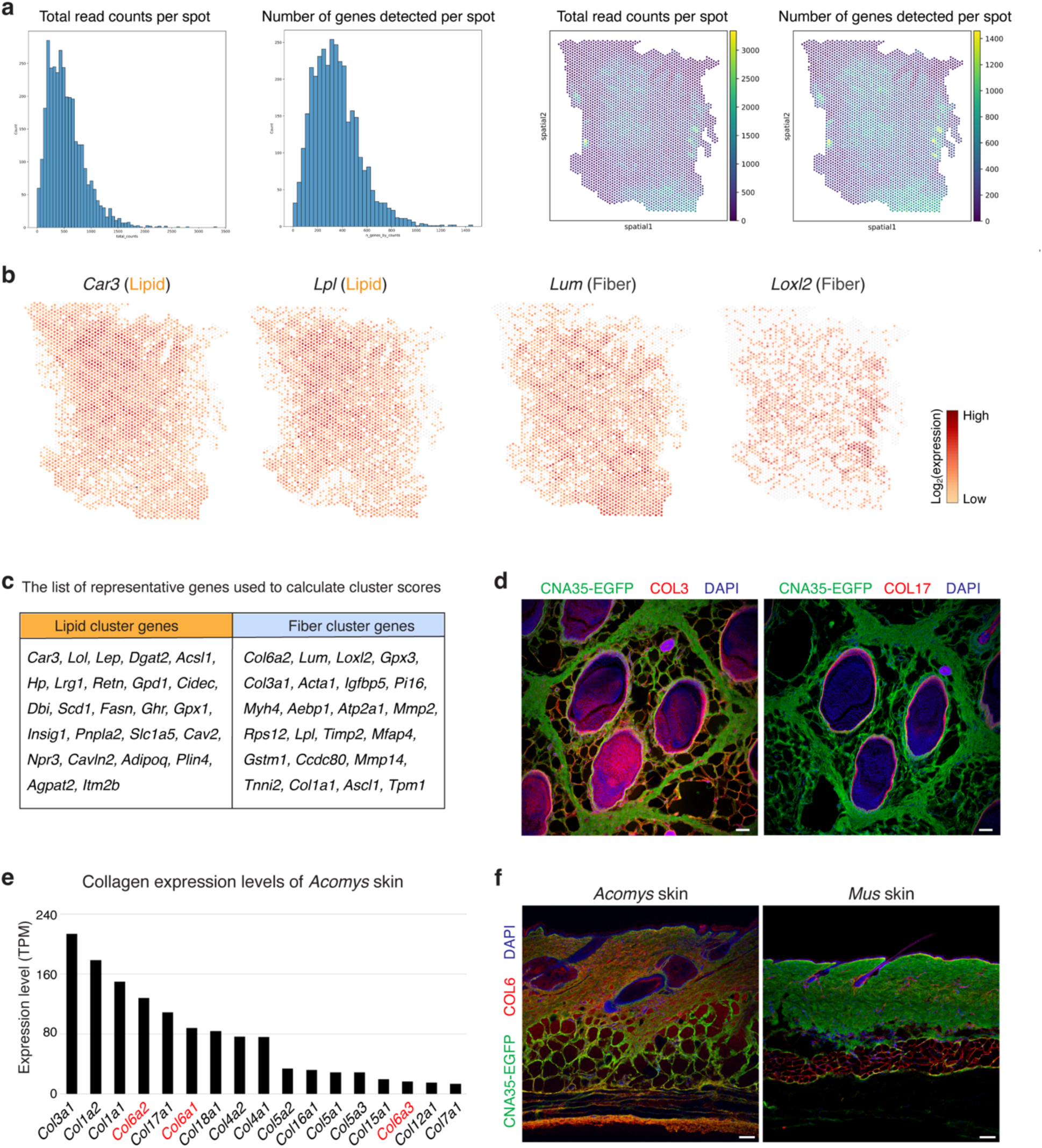
Spatial transcriptomics of *Acomys* skin. **a**, Quality control metrics displaying total read counts per spot and the number of genes detected per spot. **b**, Spatial expression patterns of “Lipid” cluster enriched genes (*Car3*, *Lpl*) and “Fiber” cluster enriched genes (*Lum*, *Loxl2*). **c**, Table summarizing the list of representative genes used to calculate cluster scores. **d**, Immunofluorescence images showing the expression pattern of COL3 and COL17 in the horizontal section of *Acomys* skin. Overall collagen structure is visualized by CNA35-EGFP. Note that COL3 is expressed in the adipose tissue, and COL17 is expressed in the basement membrane of hair follicles. None of the two collagens are expressed in the boundary of *Acomys* skin pattern. Scale bars, 100 μm. **e**, Bar graph presenting the expression levels of collagen genes in *Acomys* skin measured by bulk RNA sequencing. **f**, COL6 is expressed throughout *Acomys* skin. In the *Mus* skin, COL6 expression is restricted to hair follicles and hypodermal adipose tissue. Scale bars, 100 μm.

**Extended Data Fig. 4.**
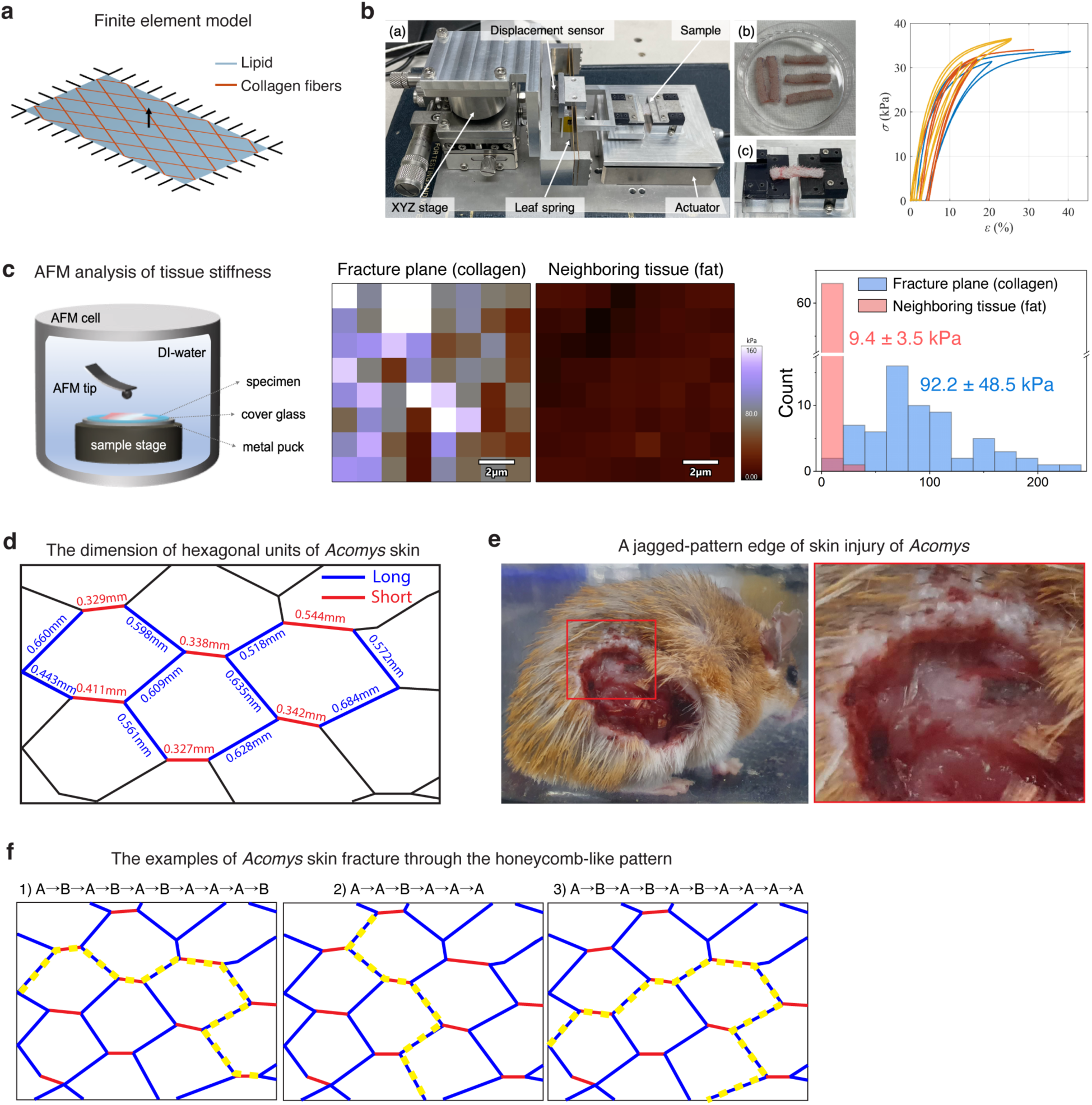
The mechanical properties of *Acomys* skin. **a**, Finite element model of *Acomys* skin with a parallelogram-shaped structure with lipids (blue) surrounded by collagen fibers (red). **b**, Tensile testing to characterize the mechanical behavior of *Acomys* skin. (a) custom-built micro-tensile testing apparatus. (b) samples prepared for tensile testing. (c) sample mounted on the testing device. A plot showing three representative stress-strain curves obtained from the tensile tests. Note that the three curves distinguished by different colors represent results from separate experiments, showing little variations. The elastic modulus of the material was measured as 644 ± 230 kPa, calculated from the loading slope of the stress-strain curve. **c**, Elastic moduli of collagen boundary and fat tissue measured by atomic force microscopy (AFM). Scale bars, 2 μm. A histogram shows the distribution of AFM measurement of collagen boundary and fat tissue. **d**, Schematic illustration of *Acomys* skin pattern, including length measurements along the boundary of each hexagon unit. **e,** Macroscopic photographs showing a jagged-pattern edge of skin injury in *Acomys*. **f**, The examples of *Acomys* skin fracture through the honeycomb arrangement.

**Extended Data Fig. 5.**
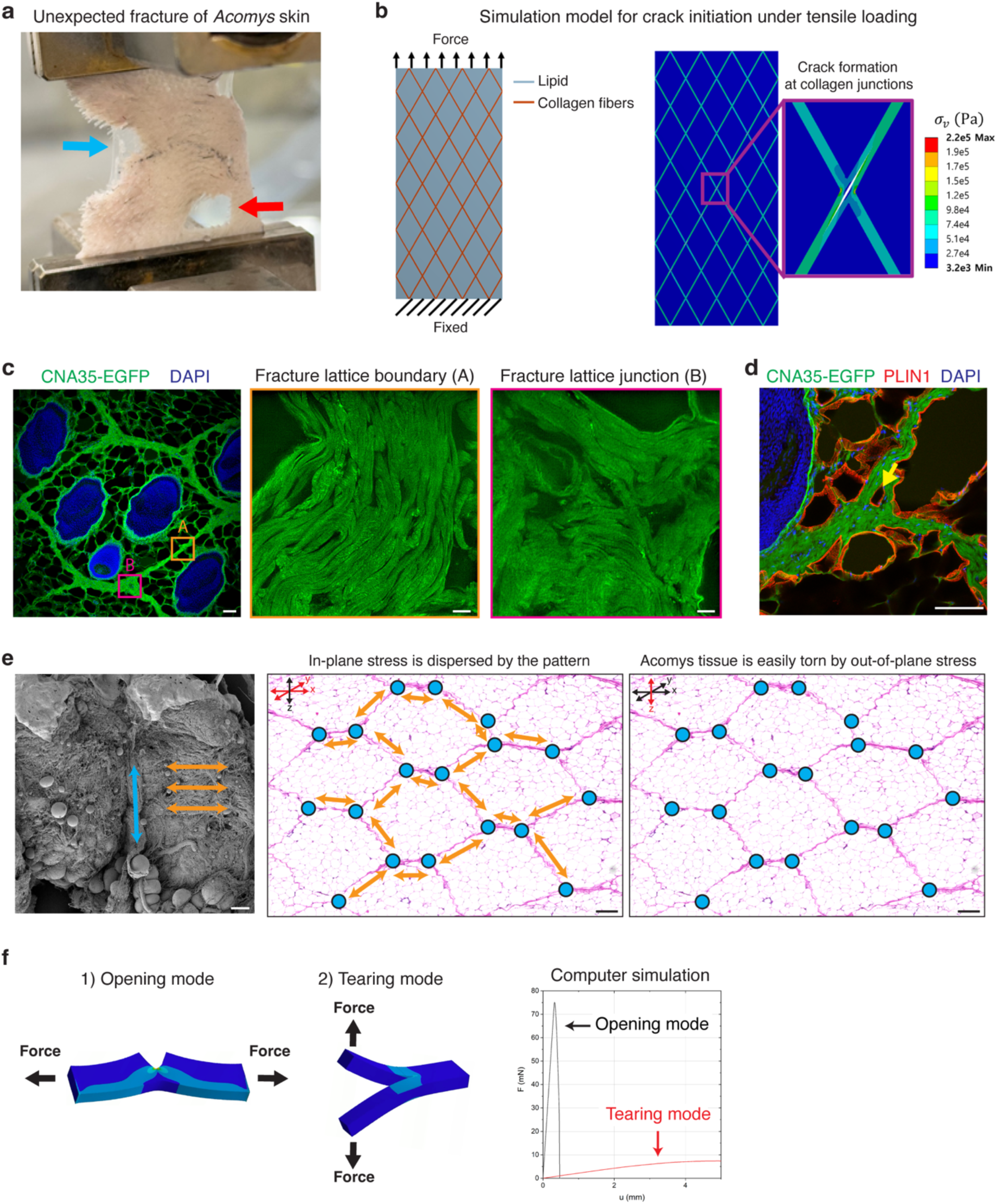
The ultrastructure of the fracture lattice in *Acomys* skin. **a**, Photograph showing unexpected fracture of *Acomys* skin during an opening mode fracture test. Blue arrow indicates a pre-defined crack, and red arrow marks unexpected fracture of *Acomys* skin. **b**, Computer simulation with a finite element model predicts the junction as a starting point of crack initiation. **c**, Structure-based super-resolution microscopy images showing CNA35-EGFP labeled collagen fibers at the boundary (yellow) and junction (red) of the fracture lattice of *Acomys* skin. Scale bars, 100 μm (left), 5 μm (middle and right). **d**, Immunofluorescence image showing a PLIN1+ adipocyte (an arrow) located in the junction of the fracture lattice. Scale bar, 100 μm. **e**, SEM and H&E images depict the directions of collagen fiber arrangement at the boundary (orange) and junction (cyan) of the fracture lattice in *Acomys* skin. In-plane stress and out-of-plane stress scenarios were shown. Scale bars, 100 μm. **f**, The results of computer simulation of opening mode and tearing mode fracture tests.

**Extended Data Fig. 6.**
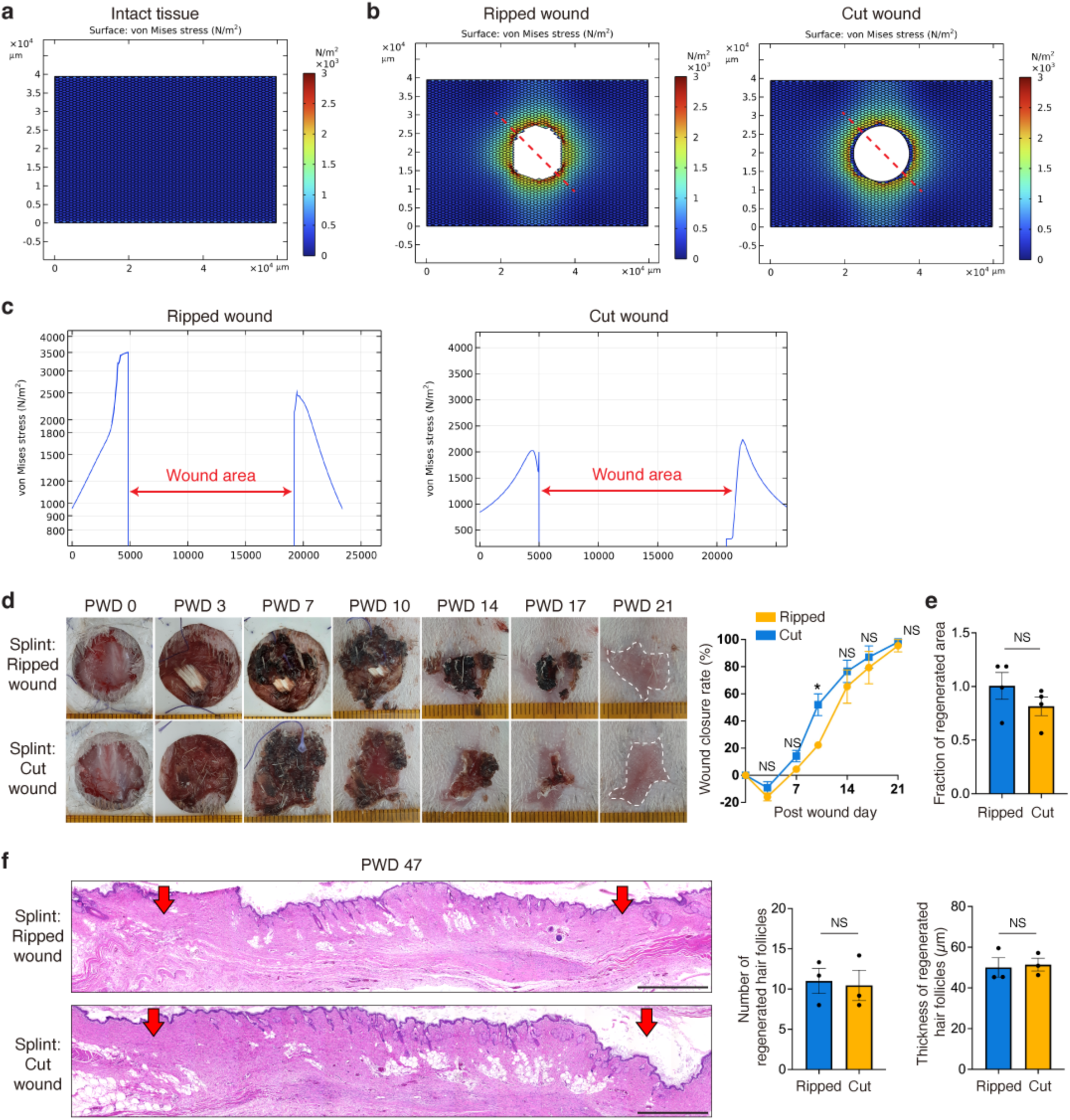
The pro-regenerative effects of ripped injury disappear by splinting. **a**, von Mises stress distributions of intact tissue containing honeycomb-like fracture lattice structure. **b**, von Mises stress distribution of ripped wound (left) and cut wound (right). **c**, Linear profiling of von Mises stress along the dotted red line in ripped wound (left) and cut wound (right). **d**, Representative photographs and a quantification graph show that wound closure rate is comparable between ripped (intergranular damage, n=3) and cut (transgranular damage, n=4) injuries with splinting. **e**, Quantification graph indicates that regenerated areas at PWD 32 are comparable between ripped (n=3) and cut (n=4) injuries. Note that the fraction of regenerated area is nearly 1, suggestive of effective inhibition of dermal contraction by splinting. **f**, H&E images and quantification graphs (n=3) show that the number and thickness of regenerated hair follicles decreased to a similar level in both ripped and cut injuries with splinting. Scale bars, 1 mm. For **d**, **e**, and **f**, statistical analysis was performed using unpaired two-tailed Student’s t-tests. Data are mean ± s.e.m.

**Extended Data Fig. 7.**
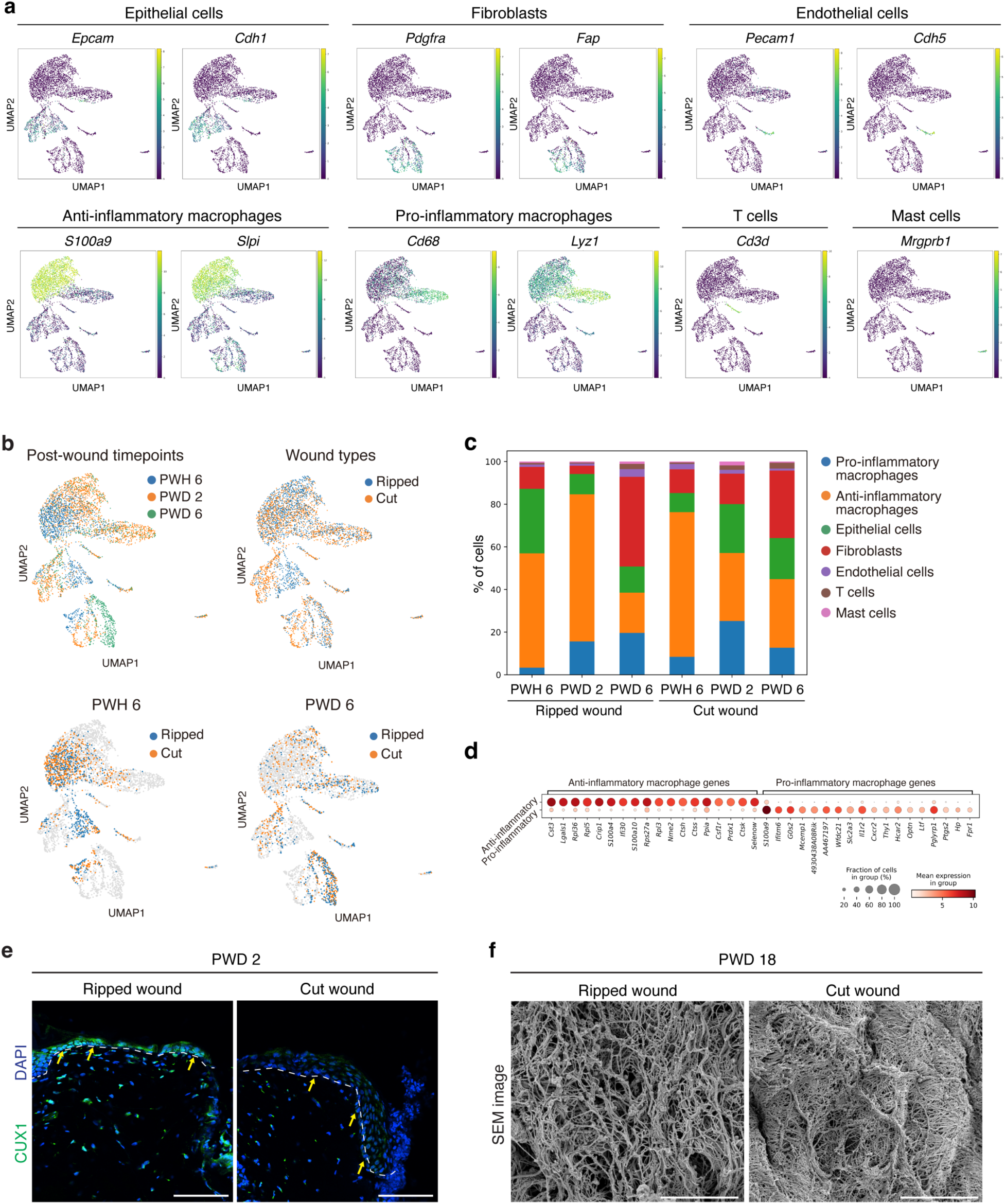
Single cell transcriptome analysis of *Acomys* skin after wounding. **a,** UMAP plots showing the expression level of signature genes for each annotated cluster of cells. **b,** UMAP plots showing single cell transcriptome analyses across wound types and post-wound timepoints. **c,** Bar plot showing the proportions of cell population across wound types and days post wounding. **d,** Dot plot showing differentially expressed genes between the clusters of pro-inflammatory macrophages and anti-inflammatory macrophages. **e,** Immunofluorescence images validating the upregulation of CUX1, a hair follicle forming transcription factor, in epidermal cells of ripped wound at PWD 2. Scale bars, 100 μm. **f,** SEM images showing that at PWD 18, cut wound exhibits excessive collagen deposition compared to ripped wound. Scale bars, 5 μm.

**Extended Data Fig. 8.**
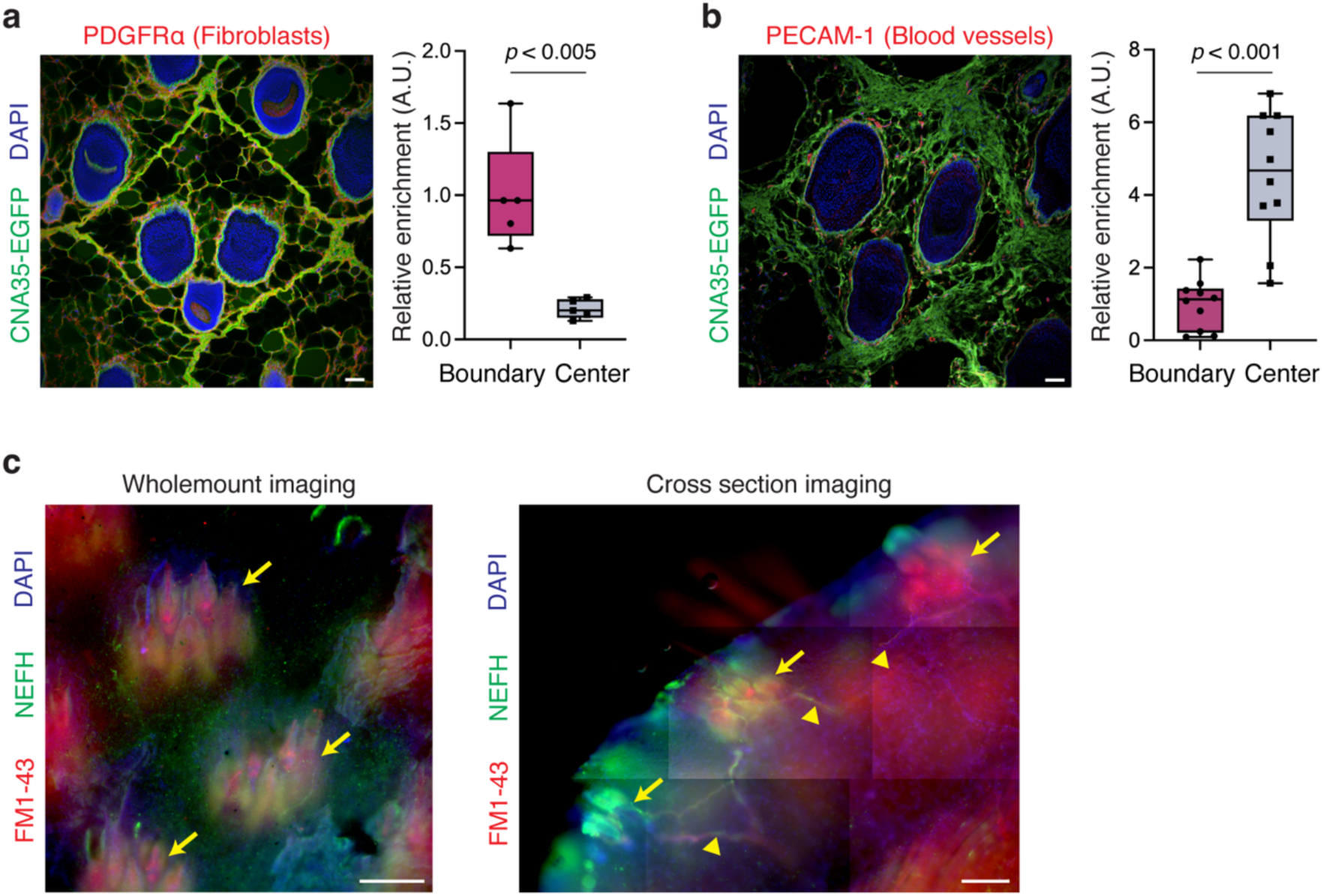
Differential distribution of skin tissue cells according to the fracture lattice. **a,** Immunofluorescence image and a quantification graph showing PDGFRα+ fibroblasts are enriched in the collagen boundary of the fracture lattice in *Acomys* skin. Scale bars, 100μm. **b**, Immunofluorescence image and a quantification graph showing PECAM-1+ blood vessels are enriched in the center regions of the fracture lattice in *Acomys* skin. Scale bars, 100μm. **c**, Wholemount and cross section skin images showing the localization of FM1-43 and NEFH-labeled putative PIEZO2^+^ neurons. Note that Acomys skin sensory neurons are concentrated to the center of the fracture lattice (yellow arrows) with few fibers extending between (yellow arrowheads). Scale bars, 200μm. For **a** and **b**, statistical analysis was performed using unpaired two-tailed Student’s t-tests. Data are mean ± s.e.m.

**Extended Data Fig. 9.**
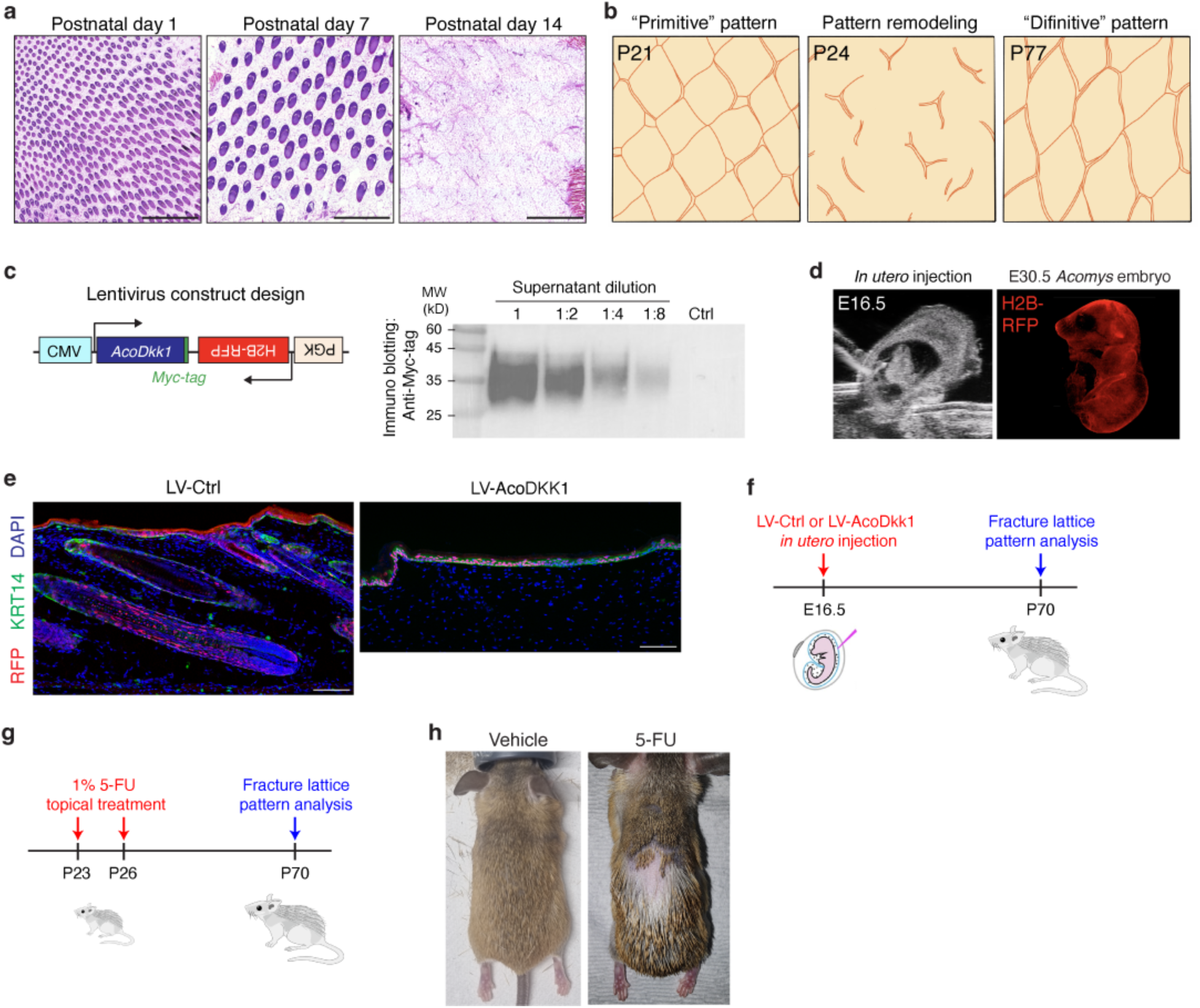
Genetic and pharmacologic manipulations for inhibiting spiny hair formation. **a**, H&E images of horizontal sections of *Acomys* back skin at different developmental stages. Note that during early development, the fracture lattice is absent. Scale bars, 1 mm. **b**, Schematic illustrations depicting the remodeling process of the fracture lattice from a primitive pattern (P21) to a definitive pattern (P77). **c**, Schematic depicting the design of lentiviral construct for overexpression *Acomys Dkk1* gene. *Acomys* DKK1 expression and secretion were confirmed by Western blot analysis using an anti-Myc-tag antibody. **d**, (left) Representative photo of ultrasound-guided lentivirus injection into the yolk sac cavity of an E16.5 *Acomys* embryo. (right) Fluorescence stereoscope image showing strong expression of H2B-RFP in an E30.5 *Acomys* embryo. **e**, Representative images showing that overexpression of AcoDKK1 effectively blocked the formation of hair follicles in *Acomys* skin. Scale bars, 200μm. **f**, Experimental timeline of AcoDKK1 overexpression and analysis of fracture lattice formation. **g**, Experimental timeline of 5-FU treatment and analysis of fracture lattice formation. **h**, Photographs showing that topical 5-FU treatment effectively inhibited spiny hair formation in *Acomys* back skin.

**Extended Data Fig. 10.**
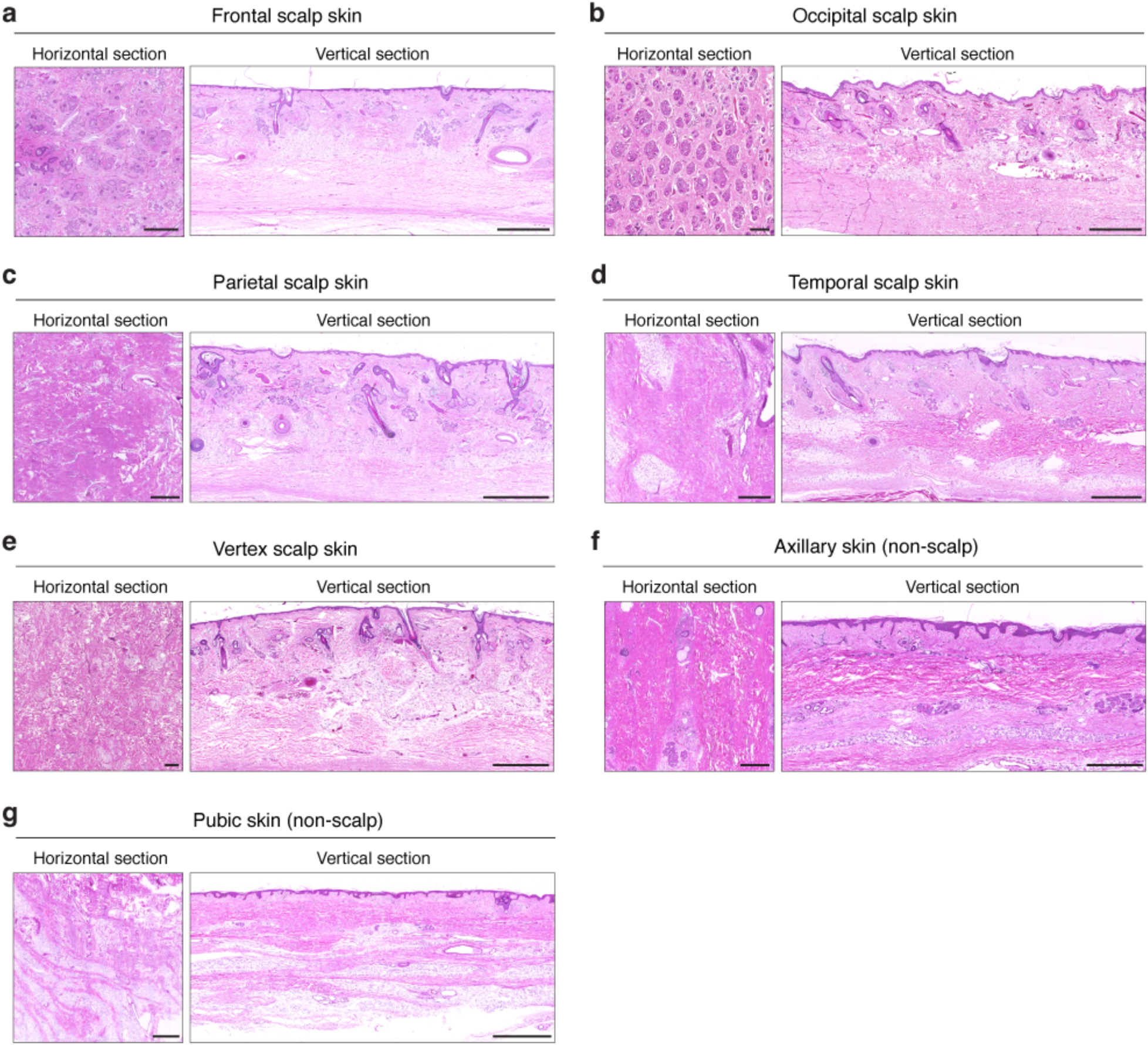
Horizontal H&E staining analyses of human scalp and non-scalp skin. **a-e**, H&E images of horizontal and vertical sections of human scalp skin. Frontal scalp skin (**a**), occipital scalp skin (**b**), parietal scalp skin (**c**), temporal scalp skin (**d**), and vertex scalp skin (**e**) were analyzed. Scale bars, 1 mm. **f,g,** H&E images of horizontal and vertical sections of human non-scalp skin. Axillary skin (**f**) and pubic skin (**g**) were analyzed. Scale bars, 1 mm.

**Extended Data Fig. 11.**
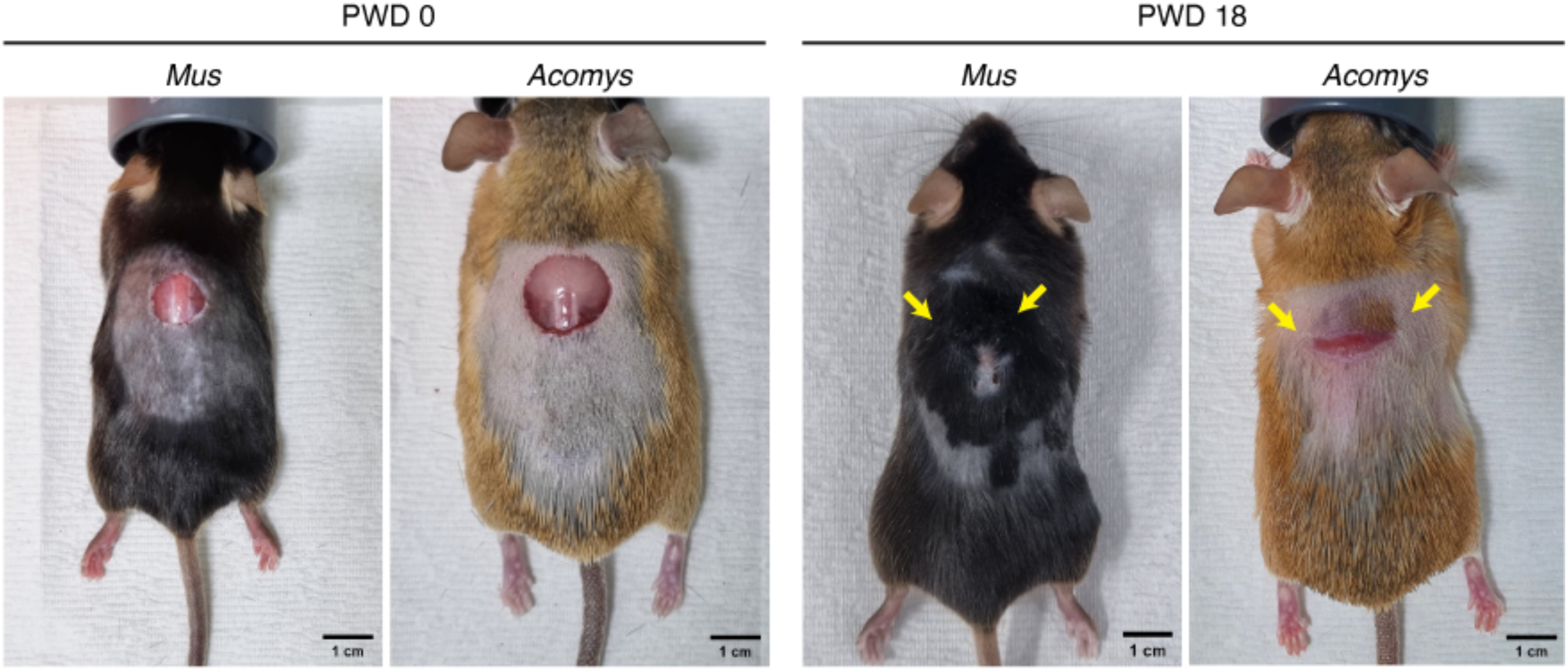
The fracture lattice potentially prevents the spread of injury signals in *Acomys* skin. Representative photographs show the wound and neighboring tissues of *Mus* and *Acomys* skin at PWD 0 and PWD 18. Note that in *Mus* skin, neighboring tissues exhibit active hair cycle induction, while those in *Acomys* remain in the resting phase. Scale bars, 1 cm.

**Extended Data Fig. 12.**
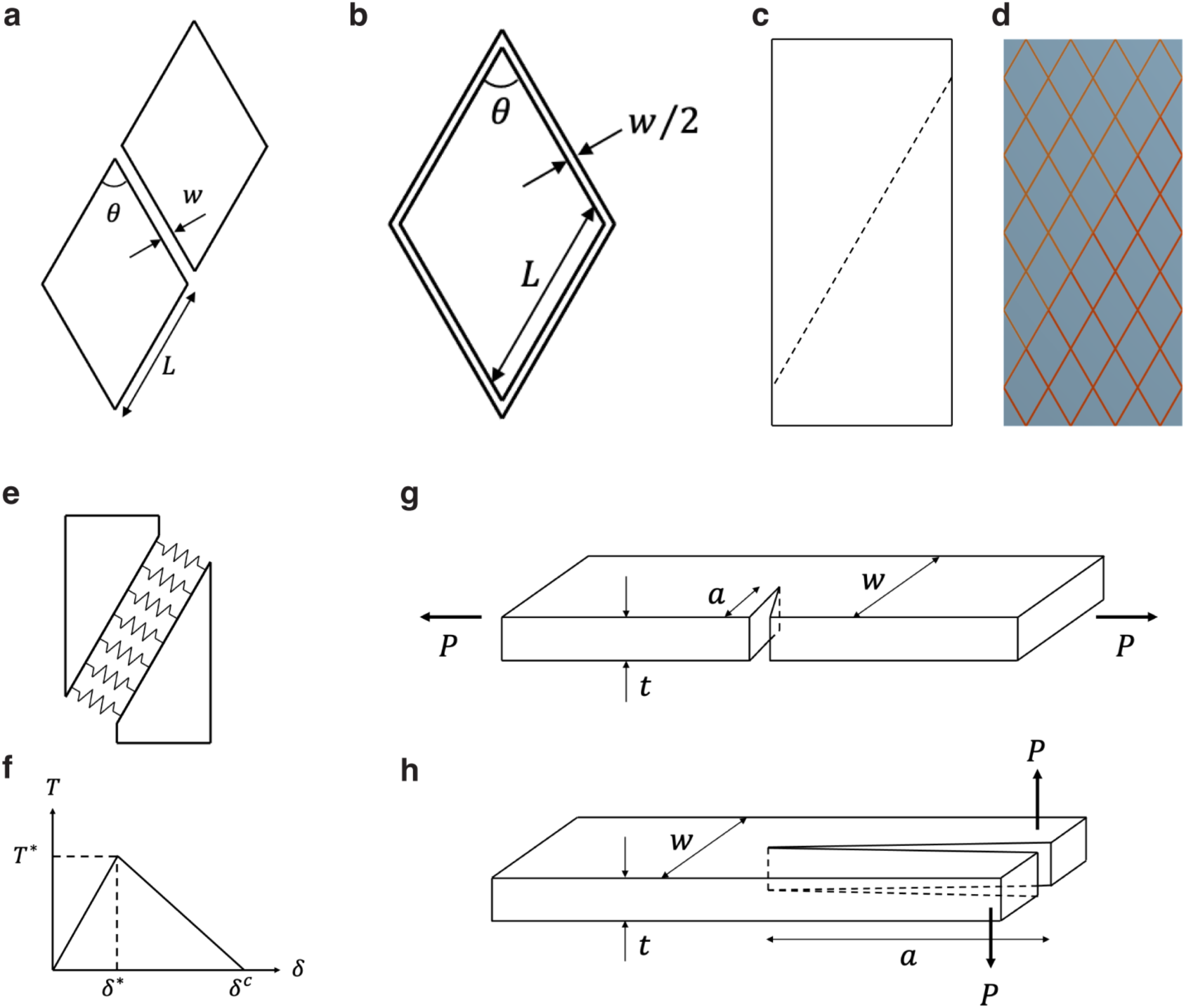
Simplified structure of *Acomys* skin and a model for evaluating resistance to crack propagation. **a,b,** Two lipid clusters (**a**) and a unit cell (**b**). *L*, the length of parallelograms. *W*, the width of lipid. θ, the internal angle of the parallelogram. **c,** The crack path indicated by a dashed line. **d,** Finite element model. **e,** Concept of the cohesive zone model. **f,** A traction-separation curve. **g,h,** Pure opening (**g**) and pure tearing (**h**) fracture. *t*, thickness. *W*, width. *a*, initial crack length. *P*, applied load.

**Supplementary Video 1.** Pinch load fracture tests in *Acomys* and *Mus* skin

**Supplementary Video 2.** Whole-mount staining of *Acomys* skin labeled by CNA35-EGFP

**Supplementary Video 3.** Pinch load simulation with and without *Acomys* skin pattern

**Supplementary Video 4.** Crack initiation simulation

**Supplementary Video 5.** Opening mode fracture tests in *Acomys* and *Mus* skin

**Supplementary Video 6.** Tearing mode fracture tests in *Acomys* and *Mus* skin

**Supplementary Video 7.** Crack propagation simulation in opening and tearing modes

## Supplementary Notes

### Elastic moduli

The elastic modulus is a fundamental material property that governs mechanical deformation, representing a material’s resistance to deformation under applied stress. As illustrated in Fig. 2C, the skin of *Acomys* exhibits a unique structure, with collagen fibers encasing lipid clusters. To accurately predict the deformation behavior of *Acomys* skin, it is essential to measure the elastic moduli of both collagen and lipid. Since *Acomys* skin failure typically occurs under tensile loading, tensile experiments were conducted to characterize the tensile properties of the combined collagen-lipid system (Extended Data Fig. 4b). Additionally, atomic force microscopy (AFM)-based nanoindentation experiments were performed to determine the stiffness ratio between collagen and lipid (Extended Data Fig. 4c), allowing for the extraction of each material’s modulus.

The elastic moduli of collagen and lipid used in the simulations were determined through volume averaging, based on tensile tests (Extended Data Fig. 4b) and nanoindentation results (Extended Data Fig. 4c). The volume-averaged elastic modulus Ē is defined as:

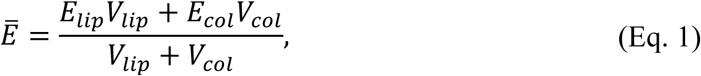

where *E* and *V* represent the elastic modulus and volume, with the subscripts “*lip*” and “*col*” referring to lipid and collagen, respectively. Nanoindentation experiments revealed that the elastic moduli of collagen and lipid exhibit an approximate ratio of *E*_*col*_⁄*E*_*lip*_ = 10 (Extended Data Fig. 4c). Using this ratio and the volume-averaged elastic modulus Ē = 644 kPa measured from tensile tests, the elastic moduli of collagen and lipid were calculated using Eq. 1, given their respective volumes.

To estimate the volume ratio, the structure of *Acomys* skin, depicted in Fig. 2C, was assumed, as illustrated in Extended Data Fig. 12a. Collagen clusters were modeled as parallelograms of length *L*, while the lipid width was defined as *W*. The unit cell of the lipid-collagen structure is shown in Extended Data Fig. 12b. Assuming *L* = 200 µm and θ = 60° for the lipid cluster and *W* = 10 µm for the collagen width, the volume ratio of lipid to collagen was calculated to be approximately 8.4. Substituting this ratio into Eq. 1, the elastic moduli of collagen and lipid were determined to be 3294 kPa and 329.4 kPa, respectively. These values were used in the finite element simulations. The Poisson’s ratio was set to 0.475, consistent with values reported for mouse skin^57^.

### Simulation under pinch loading without crack

When *Acomys* is subjected to an attack from predators, out-of-plane loads (pinch loading) are initially applied to the skin. To examine the effects of the fracture lattice structure under such conditions, a model was developed, as shown in Extended Data Fig. 4a. Shell elements were employed, with the thickness set to 2 mm to match the actual skin thickness. Displacements and rotations in all directions were constrained on the four external surfaces of the structure. A force of 0.4 N was applied to the central cell of the structure in the out-of-plane direction. Two models were considered (Fig. 3a): the first model simulates a structure composed entirely of collagen in the absence of the fracture lattice structure with an elastic modulus of *E*_*col*_ = 3294 kPa. The second model represents a structure in which collagen surrounds lipid clusters, as depicted in Fig. 3a. As shown in Fig. 3a, the presence of fracture lattice structure leads to significant stress concentration in the collagen fibers, increasing the likelihood of crack formation in the collagen.

### Crack propagation under opening mode loading

Simulations were performed to investigate the influence of the fracture lattice structure on crack initiation under tensile loading. Crack propagation was modeled using contact debonding, which does not require an initial crack. In this approach, adhesive conditions are applied to the crack surface. The crack surface was defined using an oblique crack plane, as indicated by the dashed line in Extended Data Fig. 12c. The simulations were compared to assess the influence of the lipid structure on crack propagation in the presence of the fracture lattice structures (Extended Data Fig. 12d). Contact debonding models the crack surface as a contact interface, with springs placed between the crack faces (Extended Data Fig. 12e). The spring characteristics follow a traction-separation relationship (Extended Data Fig. 12f), where *T*^∗^, δ^∗^, and δ^c^ represent the material properties governing debonding. For this study, parameters measured from meat were used: *T*^∗^ = 13.4 kPa and δ^c^ = 9.2 mm^58^.

### Opening mode (Mode I) vs Tearing mode (Mode III)

Fracture modes are typically classified as opening mode (Mode I), shearing mode (Mode II), and tearing mode (Mode III). As discussed above, when out-of-plane forces are applied to the skin, tensile loading is expected to dominate. To investigate the force required for crack propagation in thin structures, simulations were performed for both opening and tearing modes (Fig. 1E). The simulation model was designed as shown in Extended Data Fig. 12g,h, aiming to minimize mode mixity and ensure the dominance of a single fracture mode. The specimen dimensions were set as follows: total length of 20mm, thickness *t* of 2mm, and width *W* of 5mm. The initial crack length *a* was set to 2.5mm in the opening mode (Extended Data Fig. 12g) and 10mm in the tearing mode (Extended Data Fig. 12h) simulation, respectively. The crack propagation simulations were performed using an interface delamination approach, where crack propagation occurs when the energy release rate reaches a critical value. A linear fracture criterion was employed, assuming a critical energy release rate of 2 µJ⁄mm^2^ for all three fracture modes.

The crack propagation simulation videos are provided in Supplementary Video 7. The force-displacement curves derived from these simulations are presented in Fig. 4f, illustrating that crack propagation in opening mode requires substantially higher force. This observation aligns with predictions made by fracture mechanics theory. The sample geometry depicted in Extended Data Fig. 12g is commonly associated with fracture toughness predominantly governed by opening mode, and is referred to as a single edge notched tension (SENT) specimen. For this specimen, the fracture toughness can be analytically calculated as^59^:

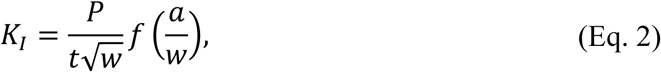

where *f* is a function of the specimen geometry. In this study, the initial crack length-to-width ratio was set to 0.5, resulting in *f*(*a*⁄*W*) of approximately 3.54^59^. Therefore, the energy release rate for the opening mode crack in Extended Data Fig. 12g is given by:

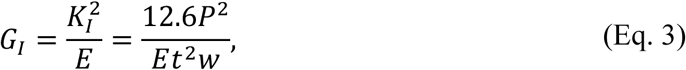

where *E* denotes the elastic modulus. For the calculation of the energy release rate in the tearing mode, the double cantilever beam model was employed. The energy release rate is given by:

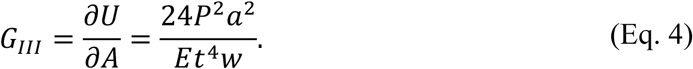

Thus, when equal forces are applied in the opening and tearing directions of the crack, the energy release rate in the tearing mode is approximately 2(*a*⁄*t*)^2^ times higher than in the opening mode. In this study, with *a*⁄*t* = 5, the energy release rate in the tearing mode is approximately 50 times greater. Therefore, if the critical energy release rates for crack propagation are identical in both modes, cracks are more likely to propagate under tearing loading. However, for short crack lengths, *G*_*I*_ may exceed *G*_*III*_. Consequently, when the crack length is less than *t*⁄√2, crack propagation occurs in the opening mode, whereas for longer cracks, propagation follows the tearing mode.

It is important to note that since the energy release rate for tearing mode was calculated using the double cantilever beam model in this section, this calculation may not be accurate for cases where the crack length-to-thickness ratio is small, as the Euler-Bernoulli beam assumption does not apply. Nevertheless, the fundamental insight that crack propagation initially occurs under opening mode, and subsequently transitions to tearing mode loading after some progression, remains valid.

## Notes

### Competing Interest Statement

The authors have declared no competing interest.

